# Circadian regulation of macromolecular complex turnover and proteome renewal

**DOI:** 10.1101/2022.09.30.509905

**Authors:** Estere Seinkmane, Anna Edmondson, Sew Y Peak-Chew, Aiwei Zeng, Nina M Rzechorzek, Nathan R James, James West, Jack Munns, David CS Wong, Andrew D Beale, John S O’Neill

## Abstract

Although costly to maintain, protein homeostasis is indispensable for normal cellular function and long-term health. In mammalian cells and tissues, daily variation in global protein synthesis has been observed, but its utility and consequences for proteome integrity are not fully understood. Using several different pulse-labelling strategies, here we gain direct insight into the relationship between protein synthesis and abundance proteome-wide. We show that protein degradation varies in-phase with protein synthesis, facilitating rhythms in turnover rather than abundance. This results in daily consolidation of proteome renewal whilst minimising changes in composition. Coupled rhythms in synthesis and turnover are especially salient to the assembly of macromolecular protein complexes, particularly the ribosome, the most abundant species of complex in the cell. Daily turnover and proteasomal degradation rhythms render cells and mice more sensitive to proteotoxic stress at specific times of day, potentially contributing to daily rhythms in the efficacy of proteasomal inhibitors against cancer. Our findings suggest that circadian rhythms function to minimise the bioenergetic cost of protein homeostasis through temporal consolidation of protein turnover.

## Introduction

Protein homeostasis, or proteostasis, refers to the dynamic process of maintaining protein abundance and functionality. It involves regulation of synthesis, folding, localisation and degradation of proteins, such that the appropriate proteins are present within the appropriate concentration range, in the correct compartment, at the right time. Multiple quality control and stress response mechanisms function to preserve proteome integrity over multiple timescales (Wolff *et al*, 2014; Harper & Bennett, 2016) whereas failure of proteostasis networks is strongly associated with impairment of cell function as well as ageing-related pathological states such as neurodegeneration (Labbadia & Morimoto, 2015; Hipp *et al*, 2019). By contrast, priming of proteostatic pathways enhances cellular resistance to proteotoxic stress (Rzechorzek *et al*, 2015).

Most aspects of mammalian cellular and organismal physiology are regulated over a circadian (about daily) timescale to anticipate the differing demands of day and night (Dibner *et al*, 2010; Atger *et al*, 2017). Whilst circadian rhythms are generated cell autonomously (Welsh *et al*, 2004; Yoo *et al*, 2004), *in vivo*, myriad cellular clocks throughout the body are synchronised with daily environmental cycles by systemic timing cues. For example, daily rhythms of feeding entrain cellular clocks through the insulin signalling pathway to stimulate PERIOD ”clock protein” production *via* activation of mammalian target of rapamycin complexes (mTORCs) (Crosby *et al*, 2019). Daily rhythms of PERIOD and mTORC activity facilitate daily rhythms of gene expression and protein synthesis. In particular, mTORC1 is a master regulator of bulk 5’-cap-dependent protein synthesis, degradation and ribosome biogenesis (Valvezan & Manning, 2019) whose activity is circadian-regulated in tissues and in cultured cells (Ramanathan *et al*, 2018; Feeney *et al*, 2016a; Stangherlin *et al*, 2021b; Mauvoisin *et al*, 2014; Jouffe *et al*, 2013; Sinturel *et al*, 2017; Cao, 2018). It is plausible that daily rhythms of mTORC activity underlie many aspects of daily physiology (Crosby *et al*, 2019; Stangherlin *et al*, 2021a; Beale *et al*, 2023b).

Most models for circadian regulation of mammalian cell function have suggested that daily rhythms in the transcription of ‘clock-controlled genes’ leads to daily rhythms in the abundance, and thus activity, of the encoded protein (Cox & Takahashi, 2019; Zhang *et al*, 2014; Andreani *et al*, 2015). However, recent -omics approaches, which measure many thousands of individual transcripts and proteins, have revealed multiple discrepancies with this abundance-based hypothesis, such as poor correlations between mRNA and encoded protein abundance (Stangherlin *et al*, 2021a). Moreover, the rather modest extent of daily changes in protein abundance (typically < 20%), and poor reproducibility between independent studies (Janich *et al*, 2015; Mauvoisin *et al*, 2014; Reddy *et al*, 2006; Robles *et al*, 2014; Mauvoisin & Gachon, 2020; Brooks *et al*, 2023), suggests that physiological variation in protein abundance is unlikely to account for large daily variations in multiple biological functions observed in tissues and cultured cells. Indeed, daily cycles of protein abundance would appear contrary to the essential requirement for maintaining proteostasis, which the major fraction of cellular energy budgets are spent to sustain (Buttgereit & Brand, 1995; Lane & Martin, 2010).

Compelling evidence for physiological daily variation in global rates of protein synthesis cannot be ignored, however (Lipton *et al*, 2015; Feeney *et al*, 2016a; Stangherlin *et al*, 2021b). Such observations are difficult to reconcile with linked observations that, excepting feeding-driven changes in mouse liver, total cellular volume and protein levels show little daily variation (Stangherlin *et al*, 2021b; Hoyle *et al*, 2017; Sinturel *et al*, 2017). To resolve these apparent discrepancies we have proposed that, in non-proliferating cells, daily changes in protein synthesis are accompanied by changes in protein degradation (Stangherlin *et al*, 2021a), resulting in daily cycles of protein turnover. This further predicts that daily rhythms in protein turnover prevail over rhythms in protein abundance to favour rhythmic proteome renewal over compositional variation. Daily turnover rhythms would be particularly beneficial for coordinated biogenesis of multiprotein complexes, since complex assembly requires individual subunits to be present stoichiometrically at the same time, in the same cellular compartment, or else be wastefully degraded (Juszkiewicz & Hegde, 2018; Taggart *et al*, 2020).

Here, we aimed to test these predictions by investigating circadian regulation of global protein synthesis and degradation, as well as macromolecular complex turnover, specifically. In so doing, we utilised bulk pulse-chase labelling to establish proof-of-principle, before developing a novel time- and cellular fraction-resolved dynamic mass spectrometry approach that provides the first direct and simultaneous measurements of protein synthesis and abundance proteome-wide. To validate our findings, we then tested for rhythmic macromolecular complex turnover directly, by quantification of nascent ribosome complex assembly through the combination of heavy uridine pulse-labelling of nascent RNA with ribosome purification. Given its importance in health and disease, we also aimed to investigate the functional consequences of rhythmic proteostasis regulation, revealing time-of-day-dependent differential sensitivity to proteotoxic stress in both cells and mice.

## Results

### Phase-coherent global rhythms in protein synthesis and degradation

We first asked whether there was any indication of cell-autonomous daily variation in protein turnover in confluent cultures of non-transformed quiescent lung fibroblasts derived from mice expressing the rhythmic luciferase reporter PER2::LUC. Using this cellular model under constant conditions, longitudinal bioluminescence recordings from parallel replicate cultures can be used to provide a robust report of cell-autonomous circadian timekeeping and establish circadian phase (Yoo *et al*, 2004; Feeney *et al*, 2016b). To measure protein degradation in parallel with synthesis, and thus assess the level of protein turnover, we first employed a traditional ^35^S-methionine/cysteine pulse-chase labelling strategy (15 min pulse, 60 min chase).

The experiment was performed over a 24h time series followed by soluble protein extraction using digitonin, which preferentially permeabilises the plasma membrane over organelle membranes. For pulse alone, ^35^S incorporation varied significantly over this period (Fig 1A, B, S1A), consistent with previous reports of rhythmic protein synthesis (Stangherlin *et al*, 2021b; Lipton *et al*, 2015; Zhuang *et al*, 2023). As expected, ∼20% of nascently synthesised proteins had been degraded after 1 hour of chase, representing rapid quality control-associated proteasomal degradation of orphan subunits as well as aberrant translation products due to premature termination and/or protein misfolding (Schubert *et al*, 2000; Wheatley *et al*, 1980; Harper & Bennett, 2016). Importantly, the proportion of degraded protein varied over time, being highest at around the same time as increased protein synthesis (Fig 1B), indicating time-of-day variation in digitonin-soluble protein turnover which cannot be solely attributed to previously reported circadian regulation of protein solubility (Stangherlin *et al*, 2021b). Rather, it suggests that global rates of protein degradation may be co-ordinated with protein synthesis rates, and may vary over the circadian cycle.

**Figure 1.**
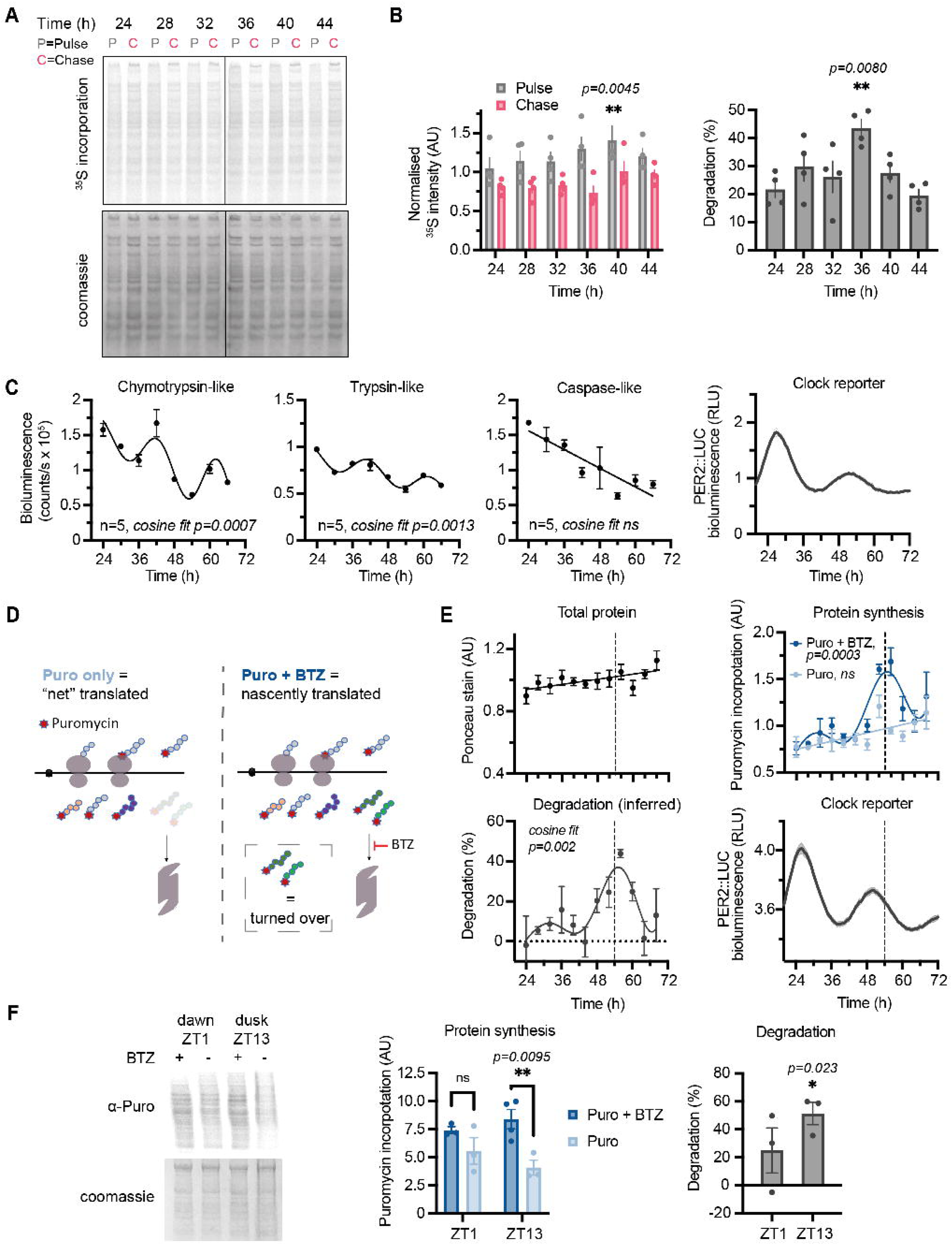
**A** A representative phosphor screen exposure of SDS-PAGE gel showing ^35^S-Met/Cys incorporation in 15 min pulse (P) and 1 h chase (C) samples at different circadian times in mouse lung fibroblasts lysed with digitonin buffer. **B** Quantification of radiolabel signal in pulse and chase at the different timepoints, normalised to protein content (by Coomassie stain) in four replicates shown in Fig S1A. Statistics: two-way ANOVA with Dunnett’s multiple comparison test, comparing T24 to other timepoints. On the right, the inferred degradation within 1h of chase is plotted, calculated as 100%*(1-Chase/Pulse). Statistics: one-way ANOVA with Dunnett’s multiple comparison test, comparing T24 to other timepoints. **C** Chymotrypsin-like, trypsin-like, and caspase-like proteasome activities, measured by ProteasomeGlo cell-based assays, at different circadian times as indicated. Statistics: damped cosine wave fit compared with straight line (null hypothesis) by extra sum-of-squares F test, the statistically preferred fit is plotted and p-value displayed. Parallel PER2::LUC bioluminescence recording from replicate cell cultures (mean +/- SEM, every 30 min) is shown below, acting as phase marker. **D** Schematic representation of the optimised puromycin incorporation assay. Puromycin (Puro) is incorporated into nascent peptide chains during translation elongation. A subset of these gets degraded by the proteasome within the 30 min labelling timeframe, resulting in a measure of “net” translation. In the presence of bortezomib (BTZ), the proteasome is inhibited, so the peptides that would have been degraded are still present and can be detected. Thus, all nascently translated peptide chains can be detected. Degradation can be inferred from comparing the two conditions, and allows estimation of both translation and degradation in a single assay. **E** Quantification of total protein, protein synthesis and protein turnover from a puromycin incorporation timecourse, where at each of the 12 timepoints Puro ± BTZ was added directly to cell media, and cells lysed 30 min afterwards in digitonin buffer. Puromycin incorporation was assessed by Western blotting, and total protein from a parallel Ponceau Red stain. Change in degradation was calculated from fitted data of puromycin incorporation, relative to mean degradation level. Statistics: damped cosine wave fit compared with straight line (null hypothesis) by extra sum-of-squares F test, the statistically preferred fit is plotted and p-value displayed. Parallel PER2::LUC bioluminescence recording from replicate cell cultures (mean +/- SEM, every 30 min) is shown below, acting as phase marker (example phase comparison is shown as dashed line). **F** Puromycin incorporation *in vivo*: mice received an i.p. injection of puromycin with or without BTZ at ZT1 or ZT13 (note: zeitgeber = ‘time-giver’ and indicates hours since initial lights on). Livers were harvested 40 min afterwards and extracted with urea/thiourea buffer. Representative anti-puromycin Western blot is shown, and quantification is shown on the right, normalised for protein loading as assessed by Coomassie staining. Statistics: two-way ANOVA with Sidak’s multiple comparisons test. Four mice were used per condition, but in some cases one of the four injections were not successful i.e. no puromycin labelling was observed and so no quantification could be performed (full data in Fig S2B).

Quality control-associated degradation predominantly occurs via the ubiquitin-proteasome system (UPS) (Schubert *et al*, 2000; Wang *et al*, 2013). To directly test whether the global rate of proteasomal protein degradation is under circadian control, as suggested previously (Desvergne *et al*, 2016; Ryzhikov *et al*, 2019; Hansen *et al*, 2021), we employed biochemical in-cell assays for proteasomal activity at discrete biological times over the circadian cycle. Over two days under constant conditions we observed a significant ∼24h oscillation in proteasomal trypsin-like and chymotrypsin-like activities of the proteasome, but not caspase-like activity (Fig 1C). Moreover, we detected a significant interaction between genotype and biological time when comparing trypsin-like proteasome activity between wild type and Cryptochrome1/2-deficient cells, that lack canonical circadian transcriptional feedback repression (Fig S1B, (Wong *et al*, 2022)). Previous proteomics studies under similar conditions have revealed minimal circadian variation in proteasome subunit abundance (Wong *et al*, 2022), suggesting that proteasome activity rhythmicity, and therefore rhythms in UPS-mediated protein degradation, are regulated post-translationally (Marshall & Vierstra, 2019; Hansen *et al*, 2021).

If global rates of proteasome-mediated protein degradation vary in phase with protein synthesis over the circadian cycle, this would result in circadian organisation of nascent protein turnover. To validate this, we employed puromycin, an antibiotic which is incorporated into nascent polypeptides by both elongating and stalled ribosomes (Nathans, 1964; Semenkov *et al*, 1992) thus making these peptides amenable to immunodetection (Aviner, 2020; Goodman & Hornberger, 2013; Schmidt *et al*, 2009). Unlike ^35^S-labelled and other amino acid analogues, puromycin can be added to cell media directly, without the need to remove endogenous amino acids, thereby minimising acute perturbations. Whilst most studies have used puromycin incorporation as a proxy for translation rate, we reasoned that protein degradation should also affect the observed levels of puromycylated peptides, as these peptides are prematurely terminated and are thus identified and degraded rapidly by the ubiquitin-proteasome system (Lacsina *et al*, 2012; Szeto *et al*, 2006). Acute (30 min) puromycin treatment of cells in culture, with or without proteasomal inhibition (by bortezomib, BTZ), allowed us to measure both total nascent polypeptide production (+BTZ) and the amount of nascent polypeptides remaining when the UPS remained active (-BTZ). This allowed inference of the level of UPS-mediated degradation of puromycylated peptides within each time window, as a proxy for nascent protein turnover (Fig 1D).

As determined by anti-puromycin western blots, over two days under constant conditions, puromycin incorporation in the presence of BTZ showed significant circadian variation. In contrast, cells that were treated with puromycin alone showed no such variation, and nor did total cellular protein levels (Fig 1E, Fig S2A). These observations support the possibility that phase-coherent daily rhythms in protein degradation might act in parallel with rhythms in translation rate, such that the proportion of degraded peptides vary in synchrony with those that were translated (Fig 1E).

Protein synthesis is the most energetically expensive process that most cells undertake (Buttgereit & Brand, 1995; Lane & Martin, 2010). *In vivo*, the temporal consolidation of global translation might be expected to confer a fitness advantage by organising this energetically expensive process to coincide with the biological time of greatest (anticipated) nutrient availability. In nocturnal mice, for example, hepatic ribosome biogenesis preferentially occurs at night, during the active/feeding phase (Jouffe *et al*, 2013; Jang *et al*, 2015; Sinturel *et al*, 2017). To explore the physiological relevance of our cellular observations, we adapted the puromycin ± BTZ labelling strategy *in vivo*, to test the specific prediction that nascent protein turnover is increased during the active phase compared with the rest phase. In livers isolated from mice at opposite times of day, we observed significantly higher turnover during the night (active/feeding phase, ZT13), compared with the daytime (rest/fasting phase, ZT1) (Fig 1F, Fig S2B). We suggest that both in cells and in mouse liver *in vivo*, the daily variation in nascent protein turnover could be an expected consequence of imperfect translation, which requires ubiquitous protein quality control mechanisms to shield the proteome from defective, misfolded, or orphaned (excess subunit) proteins.

### Proteome-wide investigation of circadian protein synthesis, abundance, and turnover

Beyond protein quality control, Figure 1 invited us to consider how circadian regulation of global protein degradation might interact with rhythmic synthesis to impact proteome composition more broadly, at the level of individual proteins. In the simplest scenario, protein abundance would correlate with synthesis rate; however, rhythmic degradation might attenuate variation in the abundance of rhythmically synthesised protein, or alternatively it may generate variation in the abundance of constitutively synthesised proteins. We devised a novel proteome-wide approach to test each of these scenarios.

To measure protein production and abundance simultaneously and directly over the circadian cycle, we utilised pulsed stable isotopic labelling with amino acids in culture (pSILAC), in combination with TMT-based mass spectrometry quantification to allow multiplexed measurements. Although pSILAC is normally applied continuously and protein harvested at multiple points to measure half-life (Doherty *et al*, 2009; Schwanhäusser *et al*, 2011; Ross *et al*, 2021), here we used a repeated fixed time window for SILAC labelling to measure newly-synthesised proteins (Fig 2A, B). To enable sufficient heavy labelling for detection, a 6h time window was employed, thus measuring synthesis and abundance within each quarter of the circadian cycle.

**Figure 2.**
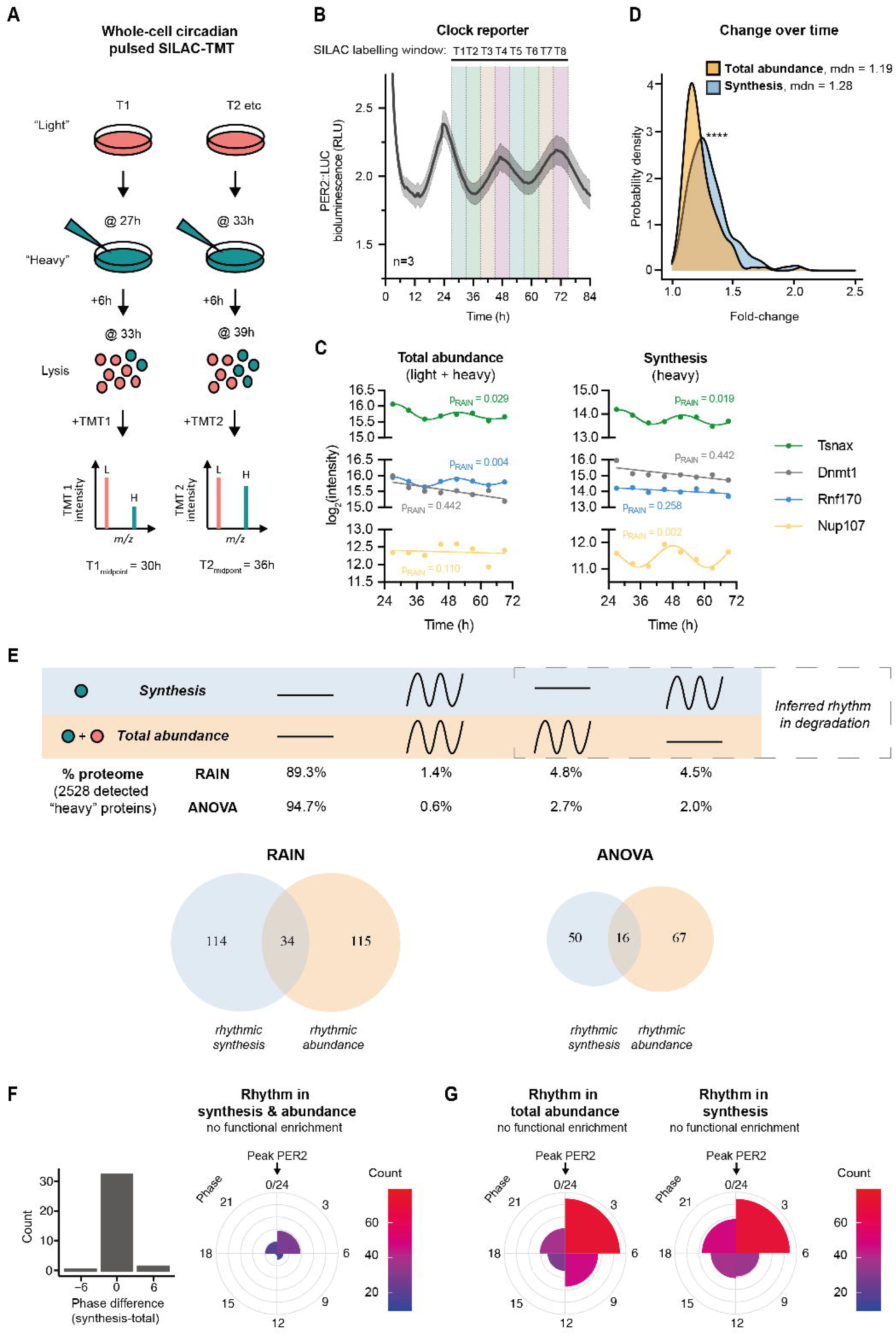
**A** Schematic of circadian pulsed SILAC-TMT experiment design. In a set of entrained fibroblasts, at each circadian timepoint “light” media (DMEM with standard L-Arg and L-Lys) is switched for “heavy” (DMEM with ^13^C_6_^15^N_4_ L-Arg and ^13^C_6_^15^N_2_ L-Lys), and cells lysed after 6h. Lysates are then digested, labelled with tandem mass tags (TMT), mixed and analysed by mass spectrometry. Data are presented aligned with the midpoint of the labelling window. **B** Parallel bioluminescence recording of PER2::LUC, acting as phase marker, overlayed with SILAC labelling windows that were used for the timecourse. **C** Representative examples of proteins (taken from Supplemental Table 1) changing rhythmically or staying constant in their total abundance (left) or synthesis (right), as measured in the pSILAC-TMT timecourse and rhythmicity defined by RAIN p<0.05. R-epresentative of rhythmic abundance and synthesis: Akap12; representative of arrhythmic abundance and synthesis: Dnmt1; representative of rhythmic abundance but arrhythmic synthesis: Rnf170; and representative of arrhythmic abundance but rhythmic synthesis: Nup107. **D** Probability density distribution of fold-change between peak and trough for proteins rhythmic in synthesis and in total abundance, representing the extent of change over time in these two sets. Statistics: Mann-Whitney test, p < 0.0001. **E** Comparison of rhythmicity between individual proteins’ synthesis and total abundance. Statistically, significant change over time was assessed by two algorithms, RAIN and ANOVA, with p<0.05 taken as rhythmicity threshold. (Top) Percentages of detected proteins falling under the four rhythmicity categories by the two algorithms are displayed; (Bottom) Venn diagrams of proteins significant for rhythms in synthesis and/or abundance. Degradation rhythms can account for cases of proteins with rhythms in synthesis but not abundance, or *vice versa*. Overall, 6264 proteins were detected; out of those at least one heavy peptide was detected for 2528 proteins (the set used for the analysis). **F** Phase distribution of proteins rhythmic (with threshold RAIN p<0.05) in both synthesis and total abundance, as well as their circadian phase difference. Gene ontology functional enrichment was tested for by GOrilla tool, in each phase separately or together, against the background of all detected proteins, and no terms were significant below the corrected p-value (FDR q-value) 0.05 cutoff. Phase 0 is set as the peak of PER2::LUC. **G** Phase distribution of proteins rhythmic (with threshold RAIN p<0.05) in their total abundance and in their synthesis. Phase 0 is set as the peak of PER2::LUC.

Over our 48h time series, we reliably detected heavy peptides for 2528 unique proteins from whole cell lysates, representing estimates of their synthesis at each time window, and compared this with their total abundance calculated from the sum of heavy and light peptides (Supplementary Table 1; examples in Fig 2C). The specific number or proportion of rhythmically synthesised and/or abundant proteins is expected to vary with detection method (Hughes *et al*, 2017; Mei *et al*, 2021) and may be susceptible to overestimation of rhythmicity. We therefore employed several methods, including less stringent RAIN and more stringent ANOVA, to compare the extent of temporal variation in protein synthesis and total abundance (Fig 2D, E).

Consistent with similar previous studies, <10% of detected proteins showed any significant variation over the circadian cycle (Fig 2E). This is also expected considering the long average half-life (∼days) of mammalian proteins (Mathieson *et al*, 2018; Schwanhäusser *et al*, 2011; Wong *et al*, 2022). Amongst those with significant temporal variation, we found that similar proportions of the proteome showed rhythms in synthesis as rhythms in abundance (Fig 2E). Of the rhythmically abundant proteins, a minority showed accompanying rhythms in synthesis, with no difference in phase (Fig 2E, F). The proportion of such proteins was more than expected by chance (p<0.0001, Fisher’s Exact Test), and their behaviour aligns with the canonical “clock-controlled gene” paradigm, in which physiological rhythms are proposed to arise through circadian variation in protein abundance, generated *via* transcriptional and translational oscillations. Strikingly however, the majority of rhythmically synthesised proteins showed no accompanying rhythm in abundance and *vice versa* (Fig 2E). Moreover, the extent of daily synthesis variation (fold change) was significantly greater than abundance (Fig 2D). These observations are consistent with our model of widespread temporal organisation of protein degradation within the circadian-regulated proteome.

Considering all detected proteins that were either rhythmically synthesised or rhythmically abundant, peaks occurred in all four quarters of the cycle (Fig 2G), but clustered in the quarter of the cycle immediately after the peak of PER2::LUC, in accordance with proteome-wide rhythms (Fig 1A, B). Gene ontology analysis did not reveal functional enrichment for any particular biological process or compartment in either group compared with background.

### Targeted investigation of circadian protein synthesis, abundance, and turnover

The experiment above was designed to combine and compare two time-resolved processes — that of circadian variation and that of protein production — and so only considered proteins with reliably detectable heavy label incorporation within a given labelling window (6h) across all timepoints. This inevitably limited and biased the proteome coverage towards abundant proteins with higher synthesis rates, irrespective of cellular compartment or function. This probably explains the absence of functional enrichment among rhythmic proteins that have been observed in other studies, as well as the lower level of overall variation in synthesis than would be expected from the bulk labelling investigations.

To gain more insight into the dynamics of circadian proteomic flux, we refined our pSILAC approach, this time focusing on proteins in complexes. Specifically, we aimed to test the hypothesis that circadian control of translation and turnover facilitates the coordinated assembly of multiprotein complexes (Taggart *et al*, 2020; O’Neill *et al*, 2020; Stangherlin *et al*, 2021a). To achieve this, we utilised and adjusted LOPIT-DC protocol (Geladaki *et al*, 2019) which was developed to separate different cellular compartments and fractions, to isolate the macromolecular complex (MMC) fraction *via* gentle cell lysis followed by sequential ultracentrifugation.

To further improve the sensitivity of our pSILAC method to circadian differences, especially those occurring among most recently synthesised proteins, we also employed a shorter pulse (1.5h). To compensate for the shorter pulse and the fractionation, which otherwise would have resulted in decreased coverage, especially of the heavy peptides, we employed further technical improvements. Namely, we added a so-called booster channel: an additional fully heavy-labelled cell sample within a TMT mixture (Klann *et al*, 2020). When the mixture is analysed by MS, heavy peptides from the booster channel increase the overall signal of all identical heavy peptides at MS1 level; at MS2 and MS3 this results in improved detection of heavy proteins in the other TMT channels of interest, and is particularly advantageous for the proteins with lower turnover that would fall below the MS1 detection limit without the booster.

With this new design, heavy peptides within the enriched MMC fraction were quantified across two days (as for the first pSILAC experiment), representing proteins synthesised within 1.5h at each timepoint (Fig 3A, B, Supplementary Table 2). Despite enriching for only one cellular compartment, the overall coverage in this experiment was similar to the previous one (6577 and 6264 proteins, respectively), due to the altered and more targeted approach; with heavy peptides detected for 2302 proteins. There was a significant circadian variation among the overall amount of heavy labelled peptides that was consistent with rhythmic production of nascent proteins, whereas the total protein level in this fraction showed no change over time (Fig S3A, B).

**Figure 3.**
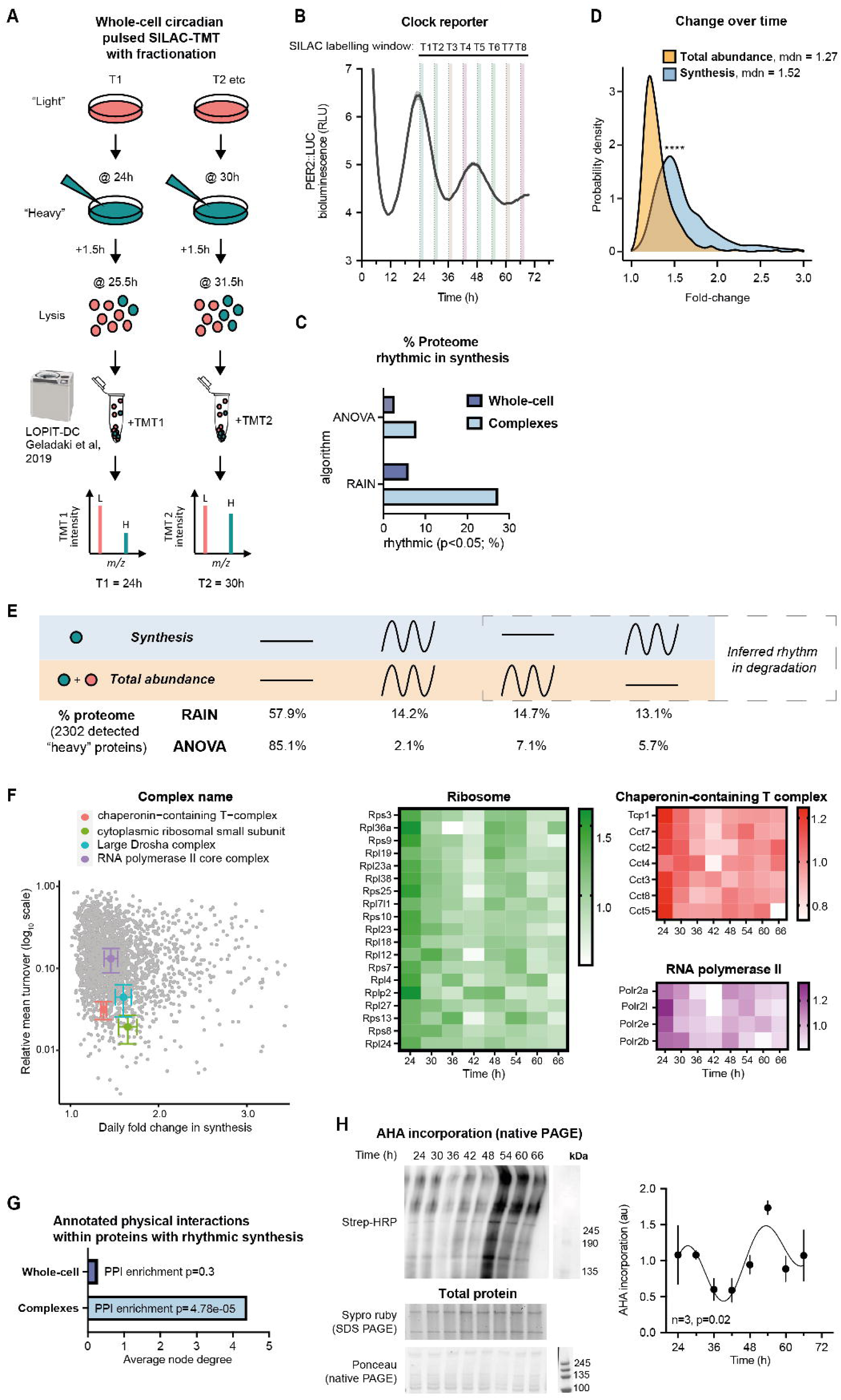
**A** Schematic of circadian pulsed SILAC-TMT with extra fractionation step. In a set of entrained fibroblasts, at each circadian timepoint “light” media (DMEM with standard L-Arg and L-Lys) is switched for “heavy” (DMEM with ^13^C_6_^15^N_4_ L-Arg and ^13^C_6_^15^N_2_ L-Lys), and cells collected after 1.5h. Samples were then subjected to sequential ultracentrifugation, a method adjusted from Geladaki *et al*. 2019 LOPIT-DC protocol. The fraction enriched for macromolecular protein complexes (MMC fraction) was labelled with tandem mass tags (TMT), mixed and analysed by mass spectrometry. **B** Parallel bioluminescence recording of PER2::LUC, acting as phase marker, overlayed with SILAC labelling windows that were used for the timecourse. **C** Comparison of percentage of detected proteins that are considered rhythmic (p<0.05) in their synthesis by the two algorithms used, between the whole-cell pSILAC experiment, presented in Fig 2, and the experiment focused on complexes, presented in this figure. **D** Probability density distribution of fold-change between peak and trough for proteins rhythmic in synthesis and in total abundance, representing the extent of change over time in these two sets. Statistics: Mann-Whitney test. **E** Comparison of rhythmicity between individual proteins’ synthesis and total abundance. Statistically, significant change over time was assessed by two algorithms, RAIN and ANOVA, with p<0.05 taken as rhythmicity threshold. Percentages of detected proteins falling under the four rhythmicity categories by the two algorithms are displayed. Overall, 6577 proteins were detected; out of those at least one heavy peptide was detected for 2302 proteins (the set used for the analysis). **F** Coordinated turnover of proteins belonging to complexes: for four selected complexes, their annotated subunits (according to a compilation of CORUM, COMPLEAT and manual annotations) were averaged in terms of their fold-change over time (x-axis), and relative turnover (proportion of heavy to total peptide intensity averaged across 8 timepoints, y axis). All proteins are displayed in the background in grey. Normalised heavy abundance, i.e. synthesis, of these proteins over time is shown on heatmaps on the right. **G** Proteins rhythmic (RAIN p<0.05) in their synthesis were analysed using STRING interaction database, filtering for high-confidence, physical interactions. Proteins with rhythmic synthesis in the complex fraction had an interconnected protein-protein interaction network, with high average node degree and significant enrichment in interactions over all detected proteins in that experiment, whereas for proteins with rhythmic synthesis in the whole-cell experiment (Fig 2) this was not the case. **H** Fibroblasts were pulsed with AHA for 1.5h at the indicated timepoints, and AHA incorporation into protein complexes and other higher molecular weight species under native conditions, using biotin as click substrate and streptavidin-HRP for detection after non-denaturing gel electrophoresis. Signal was quantified and normalised to total protein content as measured by SyproRuby (middle), and Ponceau stain of the native PAGE membrane (bottom) shows molecular weight of the loaded species. Statistics: damped cosine wave fit (plotted) preferred over straight line, extra sum-of-squares F test p-value displayed.

Using boosted fractionated pSILAC, we immediately noticed a 3-fold increase in the proportion of proteins that varied significantly over time in their synthesis as compared to the whole-cell level, regardless of algorithm used (Fig 3C). The production of rhythmically synthesised proteins in this fraction also varied over time to a far greater extent than did their abundance (∼2-fold greater variation, Fig 3D). Moreover, we found that a much higher proportion of detected proteins exhibited rhythms in both synthesis and total abundance than was observed at the whole-cell level (Fig 3E). As in whole-cell, the proportion of proteins showing rhythmic synthesis but not abundance and *vice versa*, was much greater than expected by chance (p<0.0001, Fisher’s Exact Test). By inference therefore, the proportion of proteins that are rhythmically degraded in this fraction must equal or exceed the proportion that are rhythmically synthesised. It is also noteworthy that although there were small sets of proteins that were rhythmic in both whole-cell (Figure 2) and MMC fractions (Figure 3), in both synthesis and total abundance, none of these four overlaps were higher than would have been expected by chance.

Analysis of the proteins in the MMC fraction revealed 243 annotated multiprotein complexes (from CORUM, COMPLEAT and manual annotation (Ori *et al*, 2016; Giurgiu *et al*, 2019)) to be present, including 82 complexes for which half or more annotated subunits were detected (Supplementary Table 3). It has previously been shown that protein subunits within the same complex tend to share similar turnover rates, which is thought to facilitate their co-ordinated assembly and removal (Price *et al*, 2010; Mathieson *et al*, 2018). We observed this in our data (Fig S3C) but can also add a temporal dimension: for complexes such as ribosomes, RNA polymerase, chaperonin (CCT) complex and others, the majority of component subunits not only showed similar average heavy to total protein ratios but also a similar change in synthesis over the daily cycle (Fig 3F, S3D and E). This supports the hypothesis that the assembly and turnover of macromolecular protein complexes is under circadian control.

Using an alternative approach to estimate the importance of rhythmicity for interactions of proteins within complexes, we took advantage of the STRING protein-protein interaction database (Szklarczyk *et al*, 2021). Unlike proteins with rhythmic synthesis at the whole-cell level, rhythmic proteins in this complex fraction had significantly more annotated physical interactions than would have been expected by chance given all proteins detected (Fig 3G, Fig S4). Importantly, these rhythmically synthesised protein subunits were almost all clustered within the same circadian phase (see Figure 4, discussed below).

**Figure 4.**
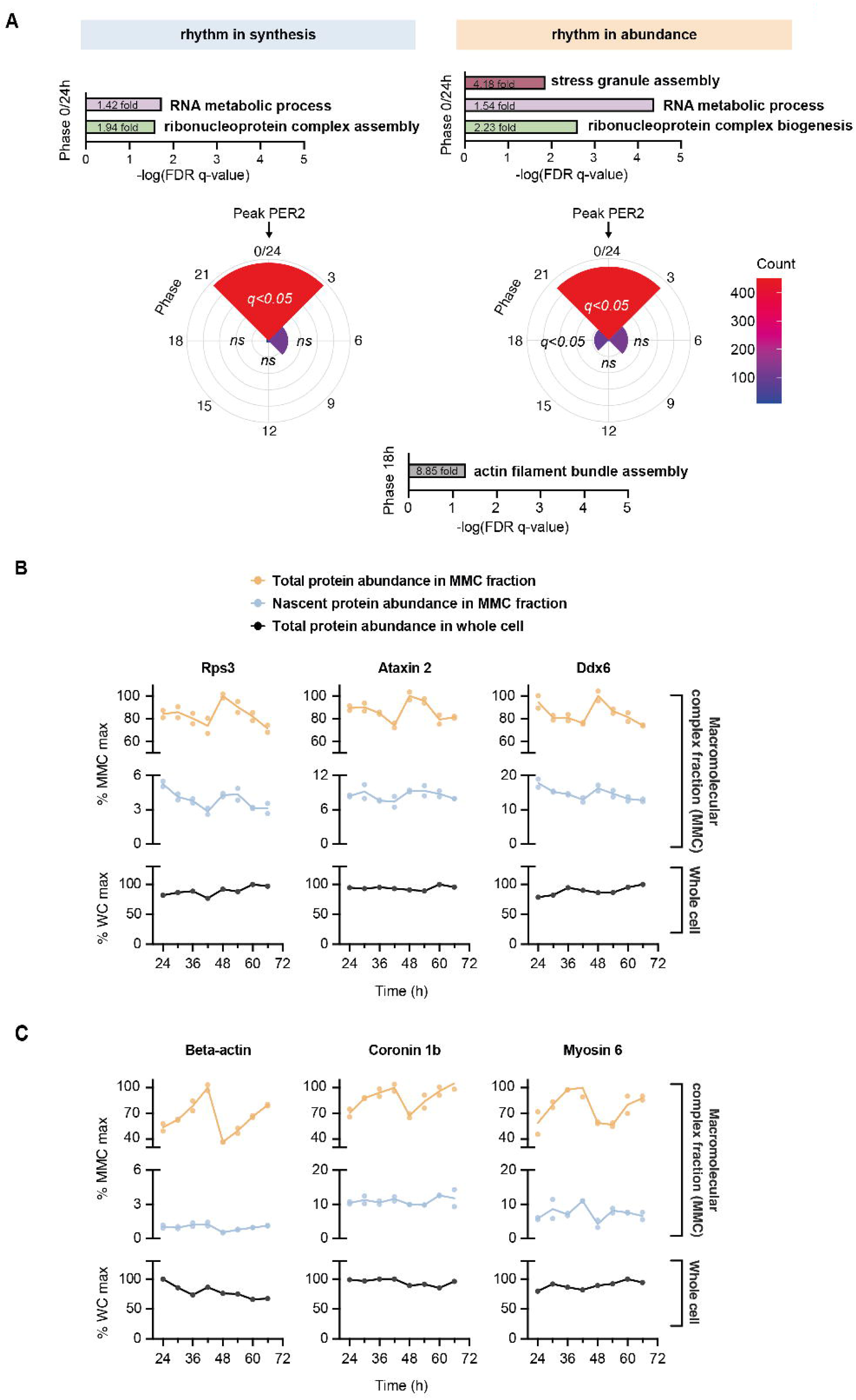
**A** Phase distribution of proteins rhythmic (RAIN p<0.05) in their total abundance and in their synthesis in the MMC fraction. Gene ontology functional enrichment for biological processes was performed using GOrilla tool, with proteins at each phase compared against all detected proteins in this experiment, and FDR q-value (multiple comparisons adjusted p-value) threshold of 0.05. For phase “0/24”, top significant non-overlapping terms are presented, alongside their significance and fold-enrichment values. In phase “18”, terms associated with actin were enriched (q<0.05), e.g. “actin filament bundle assembly” (8.85-fold enrichment). **B** Example proteins peaking at phase “0/24”, belonging to terms associated with ribonucleoprotein complex assembly and stress granule assembly. Plotted are their abundance and synthesis in the MMC fraction as well as at whole-cell level (measured independently). **C** Example proteins peaking at phase “18”, associated with actin assembly. Plotted are their abundance and synthesis in the MMC fraction as well as at whole-cell level.

To validate these observations by an orthogonal method, we pulse-labelled cells with methionine analogue L-azidohomoalanine (Dieterich *et al*, 2006). AHA is an exogenous substrate, with a lower affinity for methionyl-tRNA synthetase than methionine, whose incorporation into polypeptide chains could potentially impact the stability of labelled proteins (Ma & Yates, 2018). We therefore only used AHA to assess nascent complex synthesis, rather than turnover. We analysed the incorporation of the newly synthesised, AHA labelled proteins into highest molecular weight protein species detected under native-PAGE conditions (Fig 3H, S3F). We observed a high amplitude daily rhythm of AHA labelling, indicating the rhythmic translation and assembly of nascent protein complexes. Taken together, these results show that daily rhythms in synthesis and degradation may be particularly pertinent for subunits of macromolecular protein complexes.

### Temporal consolidation of biological functions

Within the MMC fraction, we found that the vast majority of rhythmically synthesised proteins showed highest synthesis at the same biological time, shortly after the peak of the PER2::LUC circadian bioluminescence reporter. Gene ontology analysis of these proteins (compared with a background list of all proteins detected in this fraction) revealed a clear enrichment for RNA binding proteins and terms associated with ribosome assembly (Fig 4A). At the same circadian phase, rhythmically abundant proteins were similarly enriched for terms relating to RNA binding and ribonucleoprotein biogenesis, as well as many proteins associated with stress granule assembly, such as ataxin-2 and many DDX family members (Fig 4A,B). Ribosomes and stress granules themselves control protein synthesis and regulate each other, so it is challenging to ascribe causal relationships between the two (Buchan & Parker, 2009; Riggs *et al*, 2020; Delarue *et al*, 2018) but our analysis clearly suggests a cell-autonomous surge of ribosome biogenesis.

Rhythmically synthesised/abundant proteins belonging to classes associated with ribonucleoproteins did not exhibit commensurate total abundance oscillations at the whole-cell level (Fig 2, Fig 4B, Supplementary Table 1; (Wong *et al*, 2022; Hoyle *et al*, 2017)), and this might indicate that some abundance variation in the MMC fraction arises from redistribution between lighter and denser fractions over the circadian cycle, consistent with circadian regulation of protein solubility and compartmentalisation described previously (Wang *et al*, 2019; Stangherlin *et al*, 2021b; Jang *et al*, 2015; Malcolm *et al*, 2019; Zhuang *et al*, 2023). Supporting this possibility, we noted a smaller group of rhythmically abundant proteins in the phase preceding ribosome biogenesis, without any accompanying change in synthesis. These proteins were enriched by 9-fold for actin and associated regulators of the actin cytoskeleton (q<0.05, Fig 4A, C). This is consistent with circadian regulation of cytoskeletal dynamics and actin polymerisation that we and others have described previously (Hoyle *et al*, 2017; Gerber *et al*, 2013). Indeed, as one of the most abundant cellular proteins, by mass alone, beta-actin accounted for 67% of the temporal compositional variation in the phase preceding ribosome biogenesis (Supplementary Table 2).

### Circadian regulation of ribosome turnover not abundance

The ribosome is by far the most abundant macromolecular complex in the cell (An & Harper, 2020) and showed clear evidence of circadian regulation of turnover but not abundance at the whole-cell level in our pSILAC proteomics. To validate this result, we took advantage of two important observations: (1) all fully assembled ribosomes incorporate ribosomal RNA (rRNA) which can be readily separated from most other cellular RNA by density gradient centrifugation; (2) pulse-labelling with heavy uridine-^15^N allows nascent RNA to be distinguished from pre-existing RNA. RNA could then be nuclease-digested, and the ratio of light to heavy uridine 51-monophosphate (UMP) quantified by mass spectrometry. By combining this stable isotope labelling with ribosome purification, we developed a novel cellular assay which we could use to identify nascently assembled ribosomes (Fig 5A). Circadian variation in the proportion of heavy UMP-containing assembled ribosomes, without an accompanying variation in total UMP abundance would directly demonstrate rhythmicity in ribosomal turnover.

**Figure 5.**
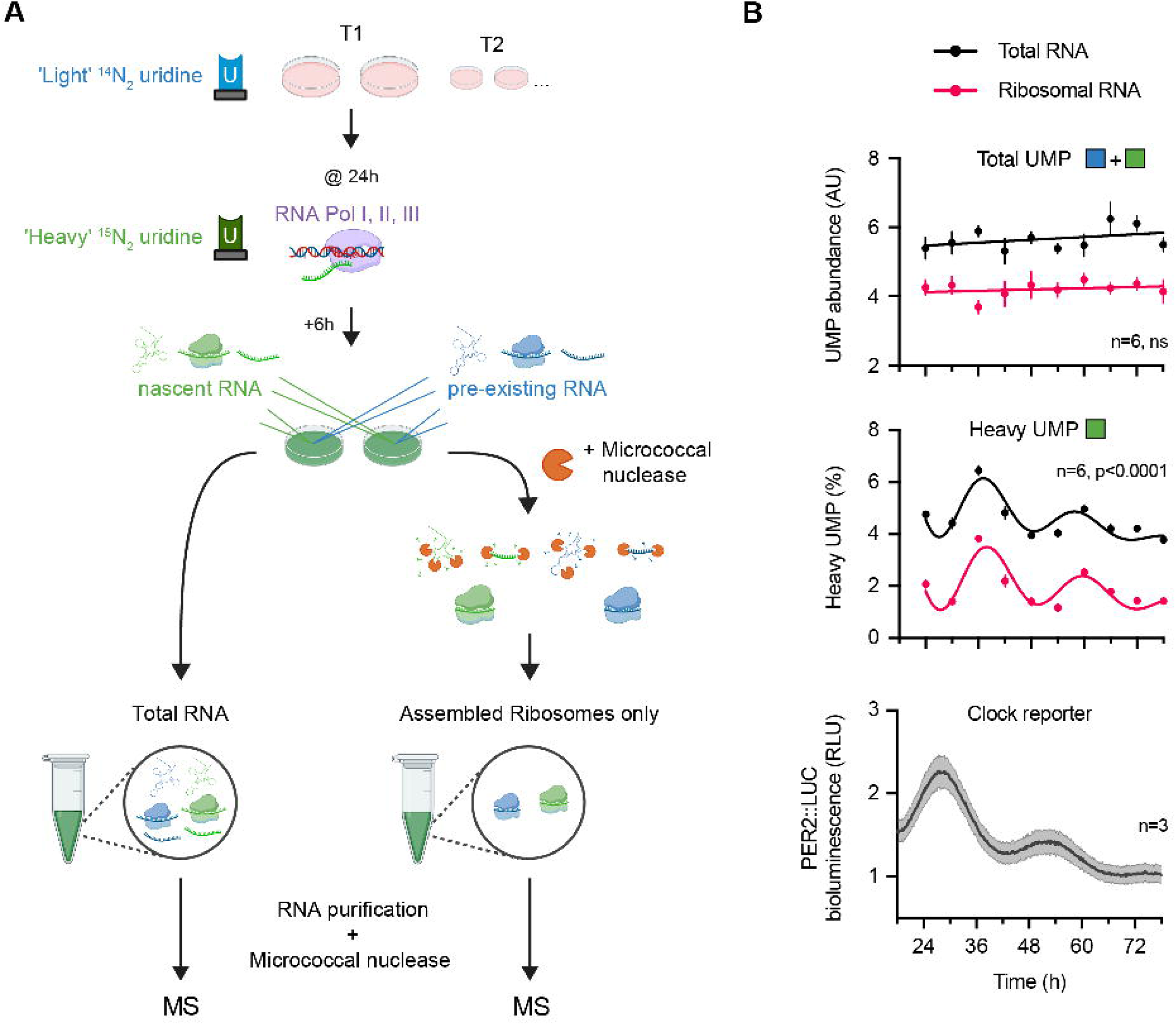
**A** Schematic of Edmondson assay for nascent rRNA labelling. In a set of entrained fibroblasts, at each circadian timepoint, 200 μM of heavy (^15^N_2_)-uridine is spiked into media for 6hr to metabolically label all nascent RNA prior to collecting cells. Total RNA or rRNA derived from assembled ribosomal complexes is extracted from cell pellets before treatment with micrococcal nuclease to completely digest RNA to free nucleotides monophosphates. The abundance of heavy and light uridine monophosphate (UMP) is then quantified by mass spectrometry. **B** Circadian regulation of ribosome turnover detected by Edmondson assay. (Top) Total UMP abundance for total cellular RNA and rRNA in assembled ribosomes. (Middle) Heavy UMP reports RNA synthesised in the preceding 6 hours, expressed as a percentage of total cellular UMP and UMP that was incorporated within assembled ribosomes, respectively. Note that heavy UMP was first corrected for natural abundance detected in unlabelled fibroblasts. (Bottom) Parallel PER2::LUC bioluminescence recording, conducted under the same experimental and timecourse conditions. Statistics: damped cosine wave fit compared with straight line (null hypothesis) by extra sum-of-squares F test, the statistically preferred fit is plotted and p-value displayed.

In line with this prediction, over a circadian time series under constant conditions, we observed an approximately 24 h oscillation in the percentage of heavy UMP detected within assembled ribosomes (Fig 5B). We found the highest rates of assembly occurred after the peak of PER2::LUC bioluminescence (Fig 5B), in line with pSILAC (Fig 3F). In addition, we observed an oscillation of heavy UMP in total RNA, likely due to rRNA which comprises >80% of total cellular RNA (Blobel & Potter, 1967; Palazzo & Lee, 2015). Importantly, total UMP within assembled ribosomes did not change significantly over time and nor did total cellular RNA (Fig 5B), providing further evidence for circadian regulation of macromolecular complex turnover rather than abundance, in line with our MMC fraction pSILAC results.

### Rhythmic response to proteotoxic stress in cells and in mice

Disruption of proteostasis and sensitivity to proteotoxic stress are strongly linked with a wide range of diseases (Wolff *et al*, 2014; Harper & Bennett, 2016; Labbadia & Morimoto, 2015; Hipp *et al*, 2019). Evidently, global protein translation, degradation and complex assembly are crucial processes for cellular proteostasis in general, so cyclic variation in these processes would be expected to have (patho)physiological consequences. Elevated levels of misfolded, unfolded, or aggregation-prone proteins perturb proteostasis and provoke proteotoxic stress responses that disrupt cellular function, leading to cell death unless resolved (Santiago *et al*, 2020; Deshaies, 2014). Informed by our observations, we predicted that circadian rhythms of global protein turnover would have functional consequences for maintenance of proteostasis. Specifically, we expected that cells would be differentially sensitive to perturbation of proteostasis induced by proteasomal inhibition using small molecules such as MG132 and BTZ, depending on time-of-day.

We first assessed the phosphorylation status of eIF2α, the primary mediator of the integrated stress response (ISR) pathway, throughout a full circadian timecourse in fibroblasts under unperturbed versus stress-induced conditions. As expected, acute proteasomal inhibition by 4h treatment with MG132 induced eIF2α phosphorylation (Jiang & Wek, 2005), but importantly this induction varied depending on time of drug treatment (Fig 6A, S5A), with highest fold-change increase observed around the predicted peak of protein turnover (shortly after the PER2::LUC peak). Phosphorylation of eIF2α leads to inhibition of canonical translation and is suggested to drive a daily decrease in bulk protein synthesis *in vivo* (Karki *et al*, 2020; Wang *et al*, 2019; Pathak *et al*, 2019). We did not observe any cell-autonomous rhythm in eIF2α phosphorylation under basal conditions (Fig S5A), and so suggest that daily p-eIF2α rhythms observed *in vivo* likely arise through the interaction between cell-autonomous mechanisms and daily cycles of systemic cues, e.g. insulin/IGF-1 signalling and body temperature rhythms driven by daily feed/fast and rest/activity cycles respectively (Crosby *et al*, 2019; Beale *et al*, 2023a).

**Figure 6.**
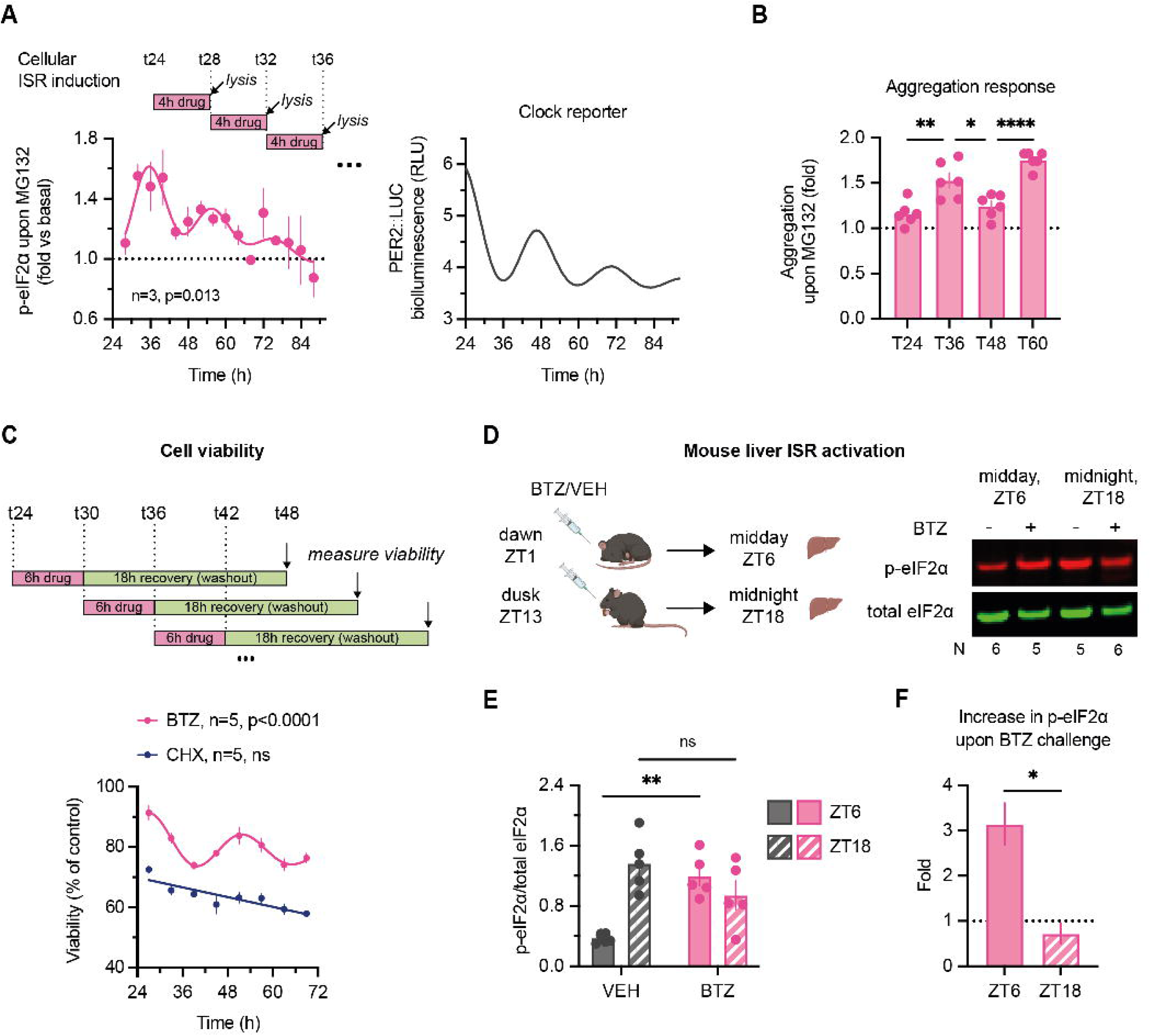
**A** (Left) Two sets of fibroblast lysates were collected every 4 h for 3 days, one untreated control and one treated with 20 µM MG132 proteasomal inhibitor for 4 h before each collection. Fold-change increase in relative phosphorylation of eIF2α (i.e. (p-eIF2α/total)_MG132_/(p-eIF2α/total)_control_) at each timepoint is plotted. MG132 is expected to induce increase in phosphorylation of eIF2α but the extent of the induction differs. Statistics: damped cosine wave fit (plotted) preferred over straight line, extra sum-of-squares F test p-value displayed. (Right) Parallel PER2::LUC bioluminescence recording, conducted under the same experimental and timecourse conditions as experiments in (A), (B) and (C). **B** Cells were treated with 20 µM MG132 for 24h starting at indicated timepoints, and aggregation relative to untreated samples measured by Proteastat aggresome kit. MG132 is expected to induce aggregation but the extent of the induction differs. Statistics: One-way ANOVA p<0.0001, stars and colours represent Dunnett’s multiple comparison of neighbouring timepoints. The difference between T24 & T48 as well as between T36 & T60 was not statistically significant. **C** At 8 timepoints throughout 2 days, fibroblasts were treated with 2.5 µM proteasomal inhibitor bortezomib (BTZ), 25 µM translation inhibitor cycloheximide (CHX), or vehicle control; after 6 h, the drugs were washed out, allowing cells to recover for further 18 h. Cellular viability after the treatments, as measured by PrestoBlue High Sensitivity assay, is expressed as a proportion of control (vehicle-treated) cells at each timepoint. Statistics: damped cosine wave fit compared with straight line (null hypothesis) by extra sum-of-squares F test, the statistically preferred fit is plotted & p-value displayed. **D** Time-of-day bortezomib (BTZ) effect *in vivo*: mice received an i.p. injection of BTZ or vehicle control (VEH) at ZT1 or ZT13, and livers were harvested 5h after the treatment. Representative Western blot is shown, blots, probed for total (green) and S51-phopshorylated (red) eIF2α. **E** Quantification of relative phosphorylation levels of eIF2α from experiment in (D). Statistics: repeated measures two-way ANOVA with Sidak’s multiple comparisons test **F** Fold-change increase in phosphorylation of eIF2α upon bortezomib injection, quantified from (E). Statistics: paired t-test.

A major detrimental consequence of proteotoxic stress is formation of insoluble intracellular protein aggregates (Albornoz *et al*, 2019; Dantuma & Lindsten, 2010). To test whether this was also time-of-day dependent, we used a molecular rotor dye that becomes fluorescent upon intercalation into quaternary structures associated with protein aggregates (Shen *et al*, 2011) (Fig S5B). As predicted, over two days, challenging cells with MG132 around the peak of protein turnover resulted in significantly more protein aggregation compared to controls than the same challenge delivered 12 h later (Fig 6B, S5C).

Sustained proteotoxic stress results in cell death (Santiago *et al*, 2020; Deshaies, 2014), and cell death induced by 6h treatment with BTZ showed a clear circadian rhythm (Fig 6C). Strikingly, we found roughly twice as much cell death occurred for proteasomal inhibition at the peak of protein turnover compared with its nadir (Fig 6C, S5D). In contrast, translational inhibition with cycloheximide revealed no such temporal variation. Together, these data support our predictions, wherein proteasomal inhibition at peak times of translation and protein turnover exacerbates proteotoxic stress, protein aggregation, and cell death because the burden on protein quality control systems at these circadian phases is already high.

BTZ and its derivatives are used clinically to treat several types of blood cancers, associated with a multitude of side effects due to the proteasome’s essential function in all cells (Deshaies, 2014; Manasanch & Orlowski, 2017; Zhang *et al*, 2020). In light of the daily variation in protein turnover we observed in mouse liver (Fig 1F), we hypothesised that time-of-day sensitivity to BTZ would also be observed *in vivo*. Accordingly, we observed a stark day vs night difference in the response to BTZ treatment in mouse liver, assessed by eIF2α phosphorylation (Fig 6D-F, Fig S5E). Consistent with this, time-of-day variation in BTZ-mediated inhibition of tumour growth was recently demonstrated in a mouse tumour model study (Wagner *et al*, 2021).

## Discussion

In this work, we provide evidence for coordinated circadian regulation of protein synthesis and degradation, resulting in rhythmic protein turnover, which is particularly significant for macromolecular complexes such as the ribosome. Across all experiments in this study, we find that protein synthesis, degradation and turnover is highest during the 6-8h that follow maximal production of the clock protein PER2. This is coincident with increased glycolytic flux and respiration (Putker *et al*, 2018), increased macromolecular crowding in the cytoplasm, decreased intracellular K^+^ concentration and increased mTORC activity (Feeney *et al*, 2016a; Stangherlin *et al*, 2021b; Wong *et al*, 2022). Just as temporal consolidation of protein synthesis is thought to increase its metabolic efficiency (O’Neill *et al*, 2020), we suggest that rhythmic turnover may serve to increase the efficiency of proteostasis by minimising deleterious changes in total cellular protein content and proteome composition.

The mechanistic underpinnings for cell-autonomous circadian regulation of the translation and degradation machineries remain to be fully explored, but are likely to be driven by daily rhythms in the activity of mTORC: a key regulator of protein synthesis and degradation as well as macromolecular crowding and sequestration (Stangherlin *et al*, 2021b, 2021a; Cao, 2018; Adegoke *et al*, 2019; Ben-Sahra & Manning, 2017; Delarue *et al*, 2018). In particular, global protein synthesis rates are greatest when mTORC1 activity is highest, in tissues and cultured cells, whereas pharmacological treatments that inhibit mTORC1 activity reduce daily variation in crowding and protein synthesis rates (Feeney *et al*, 2016a; Lipton *et al*, 2015; Stangherlin *et al*, 2021b). Given our focus on proteomic flux and translation-associated protein quality control, autophagy was not directly within the scope of this study but is also mTORC-regulated and subject to daily regulation (Ma *et al*, 2011; Ryzhikov *et al*, 2019). *In vivo*, daily regulation of mTORC activity arises primarily through growth factor signalling associated with daily feed/fast cycles (Crosby *et al*, 2019; Byles *et al*, 2021). The mechanisms facilitating cell-autonomous circadian mTORC activity rhythms are incompletely understood but may include Mg.ATP availability (Feeney *et al*, 2016a) and its direct regulation by PERIOD2 (Wu *et al*, 2019). This will be an important area for future work.

Increased translation will inevitably be associated with increased production of defective translation products, such as prematurely terminated or misfolded peptides that must be rapidly cleared by ubiquitin-proteasome system-mediated degradation (Dimitrova *et al*, 2009; Wang *et al*, 2013; Gandin & Topisirovic, 2014). Proteome-wide cycling ubiquitination sites have been recently described (Hansen *et al*, 2021); here we present evidence of cell-autonomous circadian rhythms of proteasome activity and rhythmic turnover for a greater proportion of the proteome than oscillates in abundance. Accordingly, temporal coordination was found for the synthesis of heteromeric protein complexes, in particular the ribosome, the most abundant protein complex in most mammalian cells. This highlights how, even though most mammalian proteins exhibit half-lives >24h and show little daily variation in abundance, the rate at which they are replaced can be subject to circadian regulation. This may be particularly beneficial for heteromeric complex assembly. Within the MMC fraction, we observed enrichment for specific biological functions at different times of the day, e.g., ribonucleoprotein assembly vs actin polymerisation. While bulk measurements showed clear coordination on the global scale, data from whole-cell and fractionated proteomics suggest that a combination of rhythmic synthesis, degradation, crowding and sequestration acts in concert to temporally organise rhythmic macromolecular biogenesis and assembly whilst minimising changes in overall proteome composition.

More insight into the relationship between temporal organisation and proteostasis can be gained by comparing our findings with other model systems. For example, we recently found chronic proteostasis imbalance in cells and tissues deficient for the *Cry1* and *Cry2* genes, without which circadian regulation of transcription does not persist. These cells exhibit increased proteotoxic stress as well as increased circadian variation in proteome composition compared with wild-type controls (Wong *et al*, 2022). Moreover, the temporal compartmentalisation of proteome renewal processes has a clear precedent in yeast, where metabolic oscillations arise as a direct consequence of TORC-dependent cycles of protein synthesis and sequestration that are critical for preventing deleterious protein aggregation (O’Neill *et al*, 2020). In light of similar findings in the alga *Ostreococcus tauri* (Kay *et al*, 2021; Feeney *et al*, 2016a), we speculate that promoting and minimising the energetic cost of proteostasis may be an evolutionarily conserved function of circadian and related biological rhythms.

Beyond testing two key predictions in mouse liver, a limitation is that this study was restricted to quiescent primary mouse fibroblasts. In our experience, fibroblasts are a particularly powerful and predictive model for fundamental principles of cellular circadian regulation (Hoyle *et al*, 2017). Clearly though, in future it will be necessary to extend our initial findings of protein turnover *in vivo* to fully validate that daily rhythms of protein turnover and proteome renewal occur under natural conditions (daily light/dark, feed/fast, rest/activity cycles). We predict that they will be observed across multiple mature tissues, with higher amplitude than cultured cells due to amplification of cell-intrinsic processes by daily systemic cues (hormonal and body temperature rhythms). We anticipate that the relative phases of synthesis and degradation rhythms will likely differ somewhat between tissues and physiological contexts, as recently found in growing muscle for example (Kelu *et al*, 2020).

Rhythms in transcription were not addressed in this study, but as discussed above, there is a well-established discrepancy between identities and phases of rhythmic proteins and their underlying transcript levels. Regulation at the translational level has been suggested to explain these differences, although ribosomal profiling studies have noted that on average there appears to be no delay between rhythmic transcript and nascent translation (Atger *et al*, 2015; Janich *et al*, 2015; Jang *et al*, 2015). We note, however, that ribosomal profiling reports on the level and position of ribosome-mRNA association, and so does not directly measure nascent protein production. Although a good correlate when comparing steady-state conditions, ribosome profiling also does not distinguish between active and stalled ribosomes, and does not reflect all the changes in protein synthesis that occur in dynamic cellular systems or upon perturbation that globally alter proteostasis (Liu *et al*, 2017). Upon finding evidence for global changes in protein synthesis and degradation throughout the day, the development of our pulsed SILAC method was crucial for allowing us direct insight into the regulation of protein abundance. Enabled by technological improvements in peptide detection accuracy and multiplexing, this is the first report of proteins tracked both across their lifetime (production) and across the circadian cycle. Moreover, our development of a simple pulse-labelling assay for nascent ribosome assembly likely has several applications beyond circadian research.

Finally, given the extensive links between proteome imbalance and many pathological states, daily regulation of protein metabolism has implications for health and disease. Circadian disruption is already strongly associated with impaired proteostasis, though causal mechanisms are poorly understood at this time (Bolitho *et al*, 2014; Musiek *et al*, 2018; Leng *et al*, 2019; Lipton *et al*, 2017; Wong *et al*, 2022). In this study we predicted and validated that daily turnover rhythms confer daily variation on the sensitivity of cells and tissues to a clinically relevant proteasome inhibitor. This highlights how preclinical models may help to accelerate the development of (chrono)therapies, that optimise treatment outcomes by leveraging understanding of the body’s innate daily rhythms (Cederroth *et al*, 2019).

## Methods

### Cell culture and general timecourse structure

Fibroblasts originated from mice homozygous for PER2::LUCIFERASE (Yoo *et al*, 2004), isolated from lung tissue and were immortalised by serial passaging as described previously (Seluanov *et al*, 2010). For routine culture, cells were maintained at 37°C and 5% CO_2_ in Gibco™ high glucose Dulbecco’s Modified Eagle Medium (DMEM), supplemented with 100 units/ml penicillin and 100 µg/ml streptomycin, as well as 10% Hyclone™ III FetalClone™ bovine serum (GE Healthcare). When plated for experiments, cells were grown to confluence prior to the start of assaying, which ensures contact inhibition and elimination of cell division effects during the experiments (Hoyle *et al*, 2017; Ribatti, 2017).

For all the timecourse experiments, cells were subject to temperature entrainment, consisting of 12h:12h cycles of 32°C:37°C, for at least 4 days prior to the start of assaying, with media changes if required. Unless stated otherwise, the final medium change, containing 10% serum, occurred at the anticipated transition from 37°C to 32°C, as the cells were transferred to constant 37°C. This is denoted as experimental time t=0, or start of constant conditions. Sampling began at least 24h afterwards (i.e. t=24+), to avoid any transient effects of the last serum-containing medium change and temperature shift (Balsalobre *et al*, 1998; Buhr *et al*, 2010; Beale *et al*, 2023b). Parallel recording of PER2::LUCIFERASE activity were obtained using ALLIGATOR (Cairn Research) (Crosby *et al*, 2017), and luminescence quantified in Fiji/ImageJ v2.0 (Abramoff, 2007; Schindelin *et al*, 2012).

### Cell lysis and protein quantification

For timecourse experiments requiring cell lysate collections, the procedure was based on the following. Cells were washed twice with PBS, and incubated with the indicated lysis buffers: normally either digitonin buffer (0.01% digitonin, 50 mM Tris pH 7.4, 5 mM EDTA, 150 mM NaCl) for 10 min on ice or urea/thiourea buffer for 20 min at room temperature (7 M urea, 2 M thiourea, 1% sodium deoxycholate, 20 mM Tris, 5 mM TCEP), both supplemented with protease and phosphatase inhibitor tablets (Roche, 4906845001 and 04693159001) added shortly beforehand. Cells were then scraped, and lysates transferred to Eppendorf tubes, before sonication with Bioruptor sonicator (Diagenode) at 4°C, for 2-3 cycles 30 s on/30 s off. Lysates were then centrifuged at 14000 rpm for 5 min, and supernatant either flash frozen in liquid nitrogen for future use, or taken directly for further analysis. For determination of protein concentration, Pierce bicinchoninic acid assay (BCA) (Smith *et al*, 1985) was performed in microplate format according to manufacturer’s instructions, with bovine serum albumin (BSA) protein standards diluted in the same lysis buffer as experimental samples. Pierce 660 nm assay was performed instead of BCA when samples contained thiourea.

### ^35^S pulse-chase labelling

All procedures for ^35^S pulse-chase were optimised to avoid methionine starvation, serum-containing media changes, and temperature perturbations, all of which could potentially reset circadian rhythms and obscure any cell-autonomous regulation. Fibroblasts were adapted to serum-free but otherwise complete medium starting from the last 4 days of temperature entrainment. At each timepoint, the cells were pulsed with 0.1 mCi/ml ^35^S-L-methionine and ^35^S-L-cysteine mix (EasyTag™ EXPRESS35S Protein Labeling Mix, Perkin Elmer) in methionine- and cysteine-free DMEM for 15 min. For chase, the radiolabel-containing media were replaced with standard DMEM supplemented with 2 mM (10x normal concentration) of non-radiolabelled methionine and cysteine, and cells incubated for 1h. Throughout both pulse and chase the cells were maintained at 37°C. At the end of pulse and chase periods, cells were washed with ice-cold PBS and lysed in digitonin buffer (0.01% digitonin (Invitrogen), 50 mM Tris pH 7.4, 5 mM EDTA, 150 mM NaCl for 10 min on ice). Lysates were run on NuPage™ Novex™ 4-12% Bis-Tris protein gels; the gels were then stained with Coomassie SimplyBlue^TM^ SafeStain (ThermoFisher). Gels were then dried at 80°C for 45 min and exposed overnight to a storage phosphor screen (GE Healthcare, BAS-IP SR 2025), which was subsequently imaged with Typhoon FLA700 gel scanner and quantified in Fiji/ImageJ.

### Puromycin labelling

Puromycin dihydrochloride, diluted in PBS, alone or in combination with BTZ, was added directly to cells in culture medium, as 10x bolus to a final concentration of 1 µg/ml puromycin and 1 µM BTZ. Labelling proceeded for 30 min at 37°C, after which cells were lysed in a urea/thiourea buffer and puromycin detected by Western blotting.

### AHA incorporation

At each timepoint, while still maintaining cells at 37°C, complete DMEM medium was replaced with methionine-free DMEM supplemented with AHA in combination with methionine at 30:1 ratio (Bagert *et al*, 2014) – 1 mM AHA, 33 µM Met - and 1% dialysed FBS for 90 min. Cells were lysed in digitonin buffer (HEPES rather than Tris-buffered). AHA-containing proteins were conjugated to biotin by click chemistry, by adding appropriate reagents (Jena Bioscience) to the lysates, to final concentrations of 1 mM THPTA, 1 mM CuSO_4_, 2 mM Na ascorbate, and 40 µM biotin alkyne, and incubating for 1h at room temperature. Biotinylated proteins were then detected by Western blotting.

### Western blotting

Samples for denaturing polyacrylamide gel electrophoresis (SDS-PAGE) were prepared by diluting lysates with reduced NuPage™ LDS sample buffer and heating at 70°C for 10 min. Samples were run on NuPage™ Novex™ 4-12% Bis-Tris protein gels in MES buffer or on E-PAGE 8% 48-well gels (ThermoFisher). For native running conditions, NuPAGE Tris-Acetate 3% - 8% gels were used, with buffers as per manufacturer’s instructions.

For Western blotting for puromycin and AHA incorporation measurement, chemiluminescence detection was used. Proteins were transferred from the gels to nitrocellulose membranes using an iBlot system (ThermoFisher). Membranes were stained by Ponceau as control for total protein loading, then washed, blocked, and incubated with primary antibody in the blocking buffer at 4°C overnight. Anti-puromycin antibody (PMY-2A4-2 from Developmental Studies Hybridoma bank, at 1:1000) was used with 5% milk in TBST blocking buffer, and an anti-mouse HRP-conjugated secondary antibody, while AHA-biotin was detected using Strep-HRP antibody (R-1098-1 from EpiGentek at 1:2000) in 1% BSA, 0.2% Triton X100 PBS blocking buffer, with additional 10% BSA blocking step before detection. Immobilon reagents (Millipore) were used to detect chemiluminescence. Images were analysed by densitometry in ImageLab v4.1 (BioRad).

For Western blotting of total and p-eIF2α, LICOR protocols and reagents were used. Briefly, methanol-activated PVDF-FL (Immobilon) membranes were utilised for transfer, and dried for 1h before blocking. After re-activation, membranes were blocked in Intercept TBS buffer. Primary (AHO0802 from ThermoFisher, ab32157 from Abcam, both at 1:1000) and secondary (IRDye 680RD and IRDye 800CW) antibodies were diluted in Intercept TBS buffer with addition of 0.2% Tween-20. Fluorescence was detected and quantified in Odyssey® CLx Imaging system.

### Proteasome activity assays

Cell-based ProteasomeGlo™ chymotrypsin-like and trypsin-like assays (Promega) were performed according to manufacturer’s instructions (Moravec *et al*, 2009) at multiple circadian timepoints as indicated. Briefly, cells in 96-well plates and the assay reagent were equilibrated to room temperature, before reagent addition, mixing, incubation for 10 min, and luminescence measurement with Tecan Spark 10M plate reader, with integration time of 1 s per well. For analysis, bioluminescence from negative control wells (containing only culture medium and the assay reagent, but no cells) was subtracted from all the experimental conditions.

### Aggregation assays

PROTEOSTAT® Aggresome Detection kit (Enzo Life Sciences) was used for detection of protein aggregates (Shen *et al*, 2011). Cells in 96-well plates were treated with MG132 (as indicated in the figure legends), added as 10x bolus diluted in serum-free DMEM to the pre-existing culture media, and gently titurated, to avoid cellular rhythms resetting. Cells were permeabilised and stained simultaneously with PROTEOSTAT® dye and Hoechst 33342, as per kit manufacturer’s manual. Total fluorescence in blue and red channels, and representative images of individual wells were acquired using Tecan Spark Cyto plate reader.

### Viability assays

PrestoBlue™ High Sensitivity reagent (ThermoFisher), a resazurin-based dye, was used to measure cellular viability (Boncler *et al*, 2014; Xu *et al*, 2015). Cells in 96-well plates were treated with drugs or DMSO (vehicle) controls, as indicated in figure legends, added as 10xbolus diluted in serum-free DMEM on top of existing culture media. For drug washout in the timecourse experiments, cell medium was replaced with 1% serum DMEM, to allow recovery for 18 h. The assay was then performed in line with manufacturer’s guidelines: following PrestoBlue reagent addition and incubation at 37°C for 20 min, fluorescence was measured in a Tecan Spark 10M plate reader, with excitation at 550 nm and emission at 600 nm.

### General statistics

Statistical tests were performed using GraphPad Prism (v8 and v9) and R v4, and are indicated in figure legends. *P* values are either reported in figures directly, or annotated with asterisks: * *p* ≤ 0.05; ** *p*≤ 0.01, *** *p* ≤ 0.001; **** *p* ≤ 0.0001, *ns* not significant, *p*>0.05. Number of replicates are reported as *n* or *N* (for technical and biological, respectively) in the figures; error bars represent standard error (SEM) unless stated otherwise. In cases where comparison of fits was performed, determining whether the data are better described by a straight line or a cosine wave with circadian period, the following equation was used for the latter:

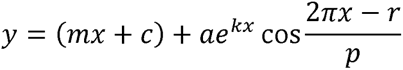

Where *m* is the baseline, *c* is the offset from 0 in y-axis, *a* is the amplitude, *k* is the damping rate, *r* is the phase, and *p* is the period, which was fixed at either 24 hours or 25 hours depending on the parallel PER2::LUC recording period.

### Proteomics data collection and analysis

#### Cell culture and sample collection for pSILAC-TMT

For pSILAC-TMT experiments, mouse lung fibroblasts were cultured in 10% dialysed FBS (dFBS). SILAC labelling was conducted in DMEM supplemented with 1% dialysed FBS and heavy-labelled amino acids instead of their light analogues, specifically 84 mg/L ^13^C_6_^15^N_4_ L-Arginine and 146 mg/L ^13^C_6_^15^N_2_ L-Lysine (Ong *et al*, 2002). In the first timecourse pSILAC experiment, sets of cells were labelled for 6h, every 6h over two days, and total cell lysates were extracted in urea/thiourea-based buffer (n=1 per timepoint). In the second timecourse experiments, labelling was done for 1.5h, every 6h over two days in duplicates, and fractionation performed as described below. For the booster channel, a fully-heavy labelled sample was used, where cells were cultured in DMEM with heavy amino acids for 5 passages (3-4 w) but otherwise processed in the same way as the timecourse samples. Fractionation was based on LOPIT-DC protocol (Geladaki *et al*, 2019). Two 15cm dishes per sample were used, cells were scraped in ice-cold PBS, centrifuged, and then lysed on ice by resuspension in a mild buffer (0.251M sucrose, 101mM HEPES pH 7.4, 21mM EDTA, 21mM magnesium acetate, protease inhibitors) and passaged through a Dounce homogeniser. Lysates were moved to thick-wall ultracentrifuge tubes (Beckman 343778 11mm/34mm) and centrifuged at 79 000 *g* for 43 min to pellet membranes and organelles. Supernatant was then centrifuged again at 120 000 *g* for 45 min. Resulting pellet was resuspended in 8 M urea 20 mM Tris buffer, and processed for mass spectrometry analysis.

### Mass spectrometry analysis

#### Protein digestion

Protein samples were reduced with 5 mM DTT at 56°C for 30 min and alkylated with 10 mM iodoacetamide in the dark at room temperature for 30 min. The samples were then diluted to 3M urea and digested with Lys-C (Promega) for 4 h at 25°C. Next, the samples were further diluted to 1.6 M urea and were digested with trypsin (Promega) overnight, at 30°C. After digestion, an equal volume of ethyl acetate was added and acidified with formic acid (FA) to a final concentration of 0.5%, mixed by shaking for 3 min and centrifuged at 15700 *g* for 2 min. The top organic layer was removed and the bottom aqueous phase was desalted using home-made C18 stage tips (3M Empore) filled with porous R3 resin (Applied Biosystems). The stage tips were equilibrated with 80% acetonitrile (MeCN) and 0.5% FA, followed by 0.5% FA. Bound peptides were eluted with 30-80% MeCN and 0.5% FA and lyophilized.

#### Tandem mass tag (TMT) labelling

Dried peptide mixtures (50 µg) from each condition were resuspended in 24 µl of 200 mM HEPES, pH 8.5. 12 µl (300 µg) TMTpro 16plex or 18plex reagent (ThermoFisher) reconstituted according to manufacturer’s instructions was added and incubated at room temperature for 1 h. The labelling reaction was then terminated by incubation with 2.2 µl 5% hydroxylamine for 30 min. The labelled peptides were pooled into a single sample and desalted using the same stage tips method as above.

#### Off-line high pH reverse-phase peptides fractionation

200 µg of the labelled peptides were separated on an off-line, high pressure liquid chromatography (HPLC). The experiment was carried out using XBridge BEH130 C18, 5 µm, 2.1 x 150 mm column (Waters), connected to an Ultimate 3000 analytical HPLC (Dionex). Peptides were separated with a gradient of 1-90% buffer A and B (A: 5% MeCN, 10 mM ammonium bicarbonate, pH8; B: MeCN, 10 mM ammonium bicarbonate, pH8, [9:1]) in 60 min at a flow rate of 250 µl/min. A total of 54 fractions were collected, which were then combined into 18 fractions and lyophilized. Dried peptides were resuspended in 1% MeCN and 0.5% FA, and desalted using C18 stage tips, ready for mass spectrometry analysis.

#### Mass spectra acquisition

The fractionated peptides were analysed by LC-MS/MS using a fully automated Ultimate 3000 RSLC nano System (ThermoFisher) fitted with a 100 μm x 2 cm PepMap100 C18 nano trap column and a 75 μm×25 cm, nanoEase M/Z HSS C18 T3 column (Waters). Peptides were separated by a non-linear gradient of 120 min, 6-38% buffer B (80% MeCN, 0.1% FA). Eluted peptides were introduced directly via a nanoFlex ion source into an Orbitrap Eclipse mass spectrometer (ThermoFisher). Data were acquired using FAIMS-Pro device, running MS3_RTS analysis, switching between two compensation voltages (CV) of -50 and -70 V.

MS1 spectra were acquired using the following settings: R = 120K; mass range = 400-1400 m/z; AGC target = 4e5; MaxIT = 50 ms. Charge states 2-5 were included and dynamic exclusion was set at 60 s. MS2 analysis were carried out with collision induced dissociation (CID) activation, ion trap detection, AGC = 1e4, MaxIT = 35 ms, CE = 34%, and isolation window = 0.7 m/z. RTS-SPS-MS3 was set up to search Uniport *Mus musculus* proteome (2021), with fixed modifications cysteine carbamidomethylation and TMTpro at the peptide N-terminal. TMTpro K, Arg10 (R +10.008), TMTpro K+K8 (K +312.221) and methionine oxidation were set as dynamic modifications. Missed cleavages were allowed, and maximum variable modifications was set at 3. In MS3 scans, the selected precursors were fragmented by high-collision dissociation (HCD), and analysed using the orbitrap with the following settings: isolation window = 0.7 m/z, NCE = 55, orbitrap resolution = 50K, scan range = 110-500 m/z, MaxIT = 200ms, and AGC = 1e5.

#### Raw MS data processing

The acquired 18 raw files from LC-MS/MS were each split into two individual spectra, one with CV = -50V and one with CV = -70V, total 36 files, using FreeStyle software (ThermoFisher). These files were then processed using MaxQuant (Cox & Mann, 2008) with the integrated Andromeda search engine (v1.6.17.0). MS/MS spectra were quantified with reporter ion MS3 from TMTpro experiments and searched against UniProt *Mus musculus* Reviewed (Nov 2020) Fasta databases. Carbamidomethylation of cysteines was set as a fixed modification, while methionine oxidation, N-terminal acetylation, Arg10 and Lys8 were set as variable modifications.

### Data analysis

After the MaxQuant search, all subsequent proteomics data processing and analysis was performed in R (v3.6.1 and v4.1.2) with R Studio v1.2. The custom scripts are available via a github repository, at https://github.com/estere-sei/circadian-pSILAC

Peptide level information from MaxQuant (evidence.txt output file) was used as a starting point. Contaminants and reverse hits were removed. Peptides were classified according to their labelling state: those that had at least one heavy arginine (Arg10) or lysine (Lys8) were classified as “heavy”, and the rest were classified as “light”. Entries for peptides with identical sequences in the same labelling state were grouped together (i.e. their reporter ion intensities across the 16 TMT channels were summed up), including peptides with other modifications such as methionine oxidation. Peptides with missing values were excluded. Total heavy label incorporation was quantified as overall proportion of summed intensities of heavy peptides over total summed intensities per TMT channel. Sample loading normalisation was performed, applying a scaling factor to equalise total summed intensity across TMT channels.

Peptides were filtered to leave only those that were detected in both heavy and light form. Peptide intensities belonging to the same leading razor protein accession were summed up to get total protein abundance value, while the sum of heavy peptides only for each protein represented the amount of synthesis. The ratio of heavy to total protein intensity averaged across the 8 timepoints was used to estimate relative turnover.

Several methods were used to assess the likelihood of significant circadian change over time in proteins’ total abundance and synthesis, including Rhythmicity Analysis Incorporating Non-parametric Methods (RAIN) (Thaben & Westermark, 2014) and ANOVA. With RAIN, the data were tested for rhythms with period length of 24 h. For ANOVA, the data were log-transformed, and two days of sampling were treated as replicates. Oscillation phase was taken from RAIN outputs, and represented circadian time of the peak of oscillation, where time 0 is equivalent to the peak of PER2::LUC from parallel recordings. The extent of change over time is expressed as fold-change, taking average ratio of peak to trough intensity values across the two days of sampling.

For protein complex membership analysis, a list was taken from Ori *et al*., 2016 which combined CORUM, COMPLEAT and manually annotated complexes and their subunits (Ori *et al*, 2016; Giurgiu *et al*, 2019; Vinayagam *et al*, 2013). Ensembl gene identifiers were converted from human to mouse by g:Profiler g:Orth tool (Raudvere *et al*, 2019), and matched with detected proteins. To asses variability of complex turnover, an analysis similar to one in Mathieson *et al*., 2018 was performed: standard deviation of the average relative turnover was calculated between proteins belonging to each detected complex, taking only complexes with more than 4 subunits, and compared to a dataset of the same size and structure but with proteins chosen randomly from all detected proteins (i.e. same number of complexes with same number of subunits as in annotated data but “subunits” chosen by random sampling).

For gene ontology functional enrichment analysis, GOrilla tool was used (Eden *et al*, 2009), comparing target protein list with all detected proteins as background, and setting FDR q-value cutoff at 0.05. REVIGO (0.4) was used to remove redundant terms. For analysis of protein-protein interactions, STRING web app was used (Szklarczyk *et al*, 2021), filtering for high-confidence physical interactions, and looking for enrichment against the background of detected proteins.

### Mouse tissue experiments

All animal work was licensed by the Home Office under the Animals (Scientific Procedures) Act 1986, with Local Ethical Review by the Medical Research Council and the University of Cambridge, UK. Throughout the experiments, wild-type C57 mice were housed in 12:12 h light:dark conditions.

For *in vivo* turnover measurements, mice received i.p. injections of either 40 µmol/kg puromycin (Ravi *et al*, 2020, 2018; Schmidt *et al*, 2009), or 40 µmol/kg puromycin in combination with 2.5 mg/kg BTZ (Apex Bio). Both solutions were sterile-filtered in PBS with 1% DMSO. Animals were culled 45 min after, in the same order as injected, and livers collected and flash frozen in liquid nitrogen. The procedure was performed twice on the same day, 1 h after the transition from dark to light (ZT1), and 1 h after the transition from light to dark (ZT13). Four age-matched male mice were used per condition.

For *in vivo* response to proteotoxic stress measurements, mice received i.p. injections of 2.5 mg/kg BTZ (Apex Bio) or vehicle control (1% DMSO in PBS, sterile-filtered). Animals were culled 5 h after, in the same order as injected, and livers collected and flash frozen in liquid nitrogen. Injections were performed twice on the same day, 1 h after the transition from dark to light (ZT1), and 1 h after the transition from light to dark (ZT13). 6 age-matched male mice were used per condition.

Tissues were homogenised in urea/thiourea lysis buffers in Precellys 24 Tissue Homogeniser (Bertin Instruments), using CK14 ceramic beads, for 3 x 15 s at 5000 rpm with 30 s breaks. Lysates were then cleared by centrifugation at 14000 rpm for 5 min, followed by protein sample preparation and Western blotting as previously described.

### Edmondson assay for nascent rRNA labelling

At each timepoint, whilst maintaining fibroblasts at constant 37°C, 200 μM isotopically heavy uridine (^15^N Cambridge Isotope Laboratories) was spiked into media for 6 h to label nascently transcribed RNA. After labelling, cells were harvested by trypsinisation, and pellets immediately flash frozen and stored at −70 °C.

For ribosome extraction, each cell pellet was resuspended in 200 μl of lysis buffer (40 mM HEPES·KOH (pH 7.5), 75 mM KOAc·HOAc, 5 mM Mg(OAc)_2_·HOAc, 1 mM CaCl_2_, 10 μM Zn(OAc)_2_·HOAc, 2 mM spermidine, 5 mM dithiothreitol, 1% v/v Triton X-100) and sonicated at 4 °C for 5 min (30 s on; 30 s off). To each lysate, 2 μl of micrococcal nuclease (Nuclease S7, Roche) was added and the lysates were incubated at 25 °C for 18 min using a thermal cycler (Techne-Prime, Cole-Parmer). Processed lysates were immediately flash-frozen in liquid nitrogen and stored at −70 °C.

Ribosomal RNA samples and cell pellets for total RNA extraction were resuspended in ‘RLT Buffer’ and RNA extracted and purified using the RNeasy Mini kit (Qiagen, 74004) according to the manufacturer’s instructions, including the on-column DNase-treatment (Qiagen, 79254). RNA was then further purified via an overnight ethanol precipitation at −20 °C, and RNA pellets resuspended in pre-heated ‘Physiological Buffer’ (50 mM HEPES·KOH (pH 7.5), 100 mM KOAc·HOAc, 20 mM Mg(OAc)_2_·HOAc). RNA was then degraded into single nucleotides by overnight room temperature incubation with micrococcal nuclease (Nuclease S7, Roche) supplemented with 1 mM CaCl_2_. The digestion reaction was then terminated by flash-freezing, and samples stored at -70°C.

### LC-MS analysis of UMP and heavy UMP in RNA lysates

10 µl of RNA lysate was diluted in 40 µl of 10 mM ammonium acetate and transferred to a 96 well skirted PCR plate (Starlab International, Hamburg, Germany) and covered with a silicone sealing mat (Axymat, Salt Lake City, Utah, USA) prior to mixed mode LC-MS analysis using an ACE Excel C18-PFP (pentafluorophenyl) column (150 × 2.1 mm, 2.0 µm, Hichrom, Reading, Berkshire, UK). Mobile phase A consisted of water with 0.1% formic acid with 10 mM ammonium formate and mobile phase B was acetonitrile with 0.1% formic acid. For gradient elution mobile phase B was held at 0% for 1.6 min. followed by a linear gradient to 30% B over 4.0 minutes, a further increase to 90% over 1 min. and a hold at 90% B for 1 min. with re-equilibration for 1.5 minutes giving a total run time of 6.5 minutes. The flow rate was 0.5 mL/min and the injection volume was 3 µL. The needle wash used was 1:1 water: acetonitrile.

For MS analysis using the Q Exactive Plus (ThermoFisher Scientific, Hemel Hempstead, Hertfordshire, UK) a full scan of 60-900 m/z was used at a resolution of 70,000 ppm in positive ion mode. The source parameters were as follows: an auxiliary gas temperature of 450°C, a capillary temperature of 275 °C, an ion spray voltage of 3.5 kV and a sheath gas, auxiliary gas and sweep gas of 55, 15 and 3 arbitrary units respectively.

Samples were analysed using Xcalibur (Thermofisher, Version 4.2) and processed and integrated using the Qual Browser and Quan Browser tools within Xcalibur to target specific analytes. Data were expressed as area ratios between areas of extracted ion chromatograms of UMP (m/z 325.0431) against 15N2 UMP (m/z 327.0372) and all identifications of compounds were carried out using reference of the accurate mass and verified using standards purchased from Sigma Aldrich.

## Acknowledgements

We thank all members of O’Neill lab, Rachel Edgar, Manu Hegde and Szymon Juszkiewicz for valuable feedback and discussions, as well as Kathryn Lilley and Holger Kramer for advice on proteomics. We also thank biomedical technical staff at Medical Research Council (MRC) Ares facility and LMB facilities for assistance. NMR was supported by the Medical Research Council (MR/S022023/1). JON was supported by the Medical Research Council (MC_UP_1201/4).

## Author contributions

ES and JON designed the study, analysed the data and wrote the manuscript with assistance from JM and ADB; SYP-C and JW performed mass spectrometry; NMR, AZ and JON performed mouse studies; ES, AE, AZ and NRJ performed cell experiments; DCSW, JM and ADB provided further experimental assistance and valuable intellectual contributions. All authors commented on the manuscript.

