## Supplementary information for "Circadian regulation of macromolecular complex turnover and proteome renewal"

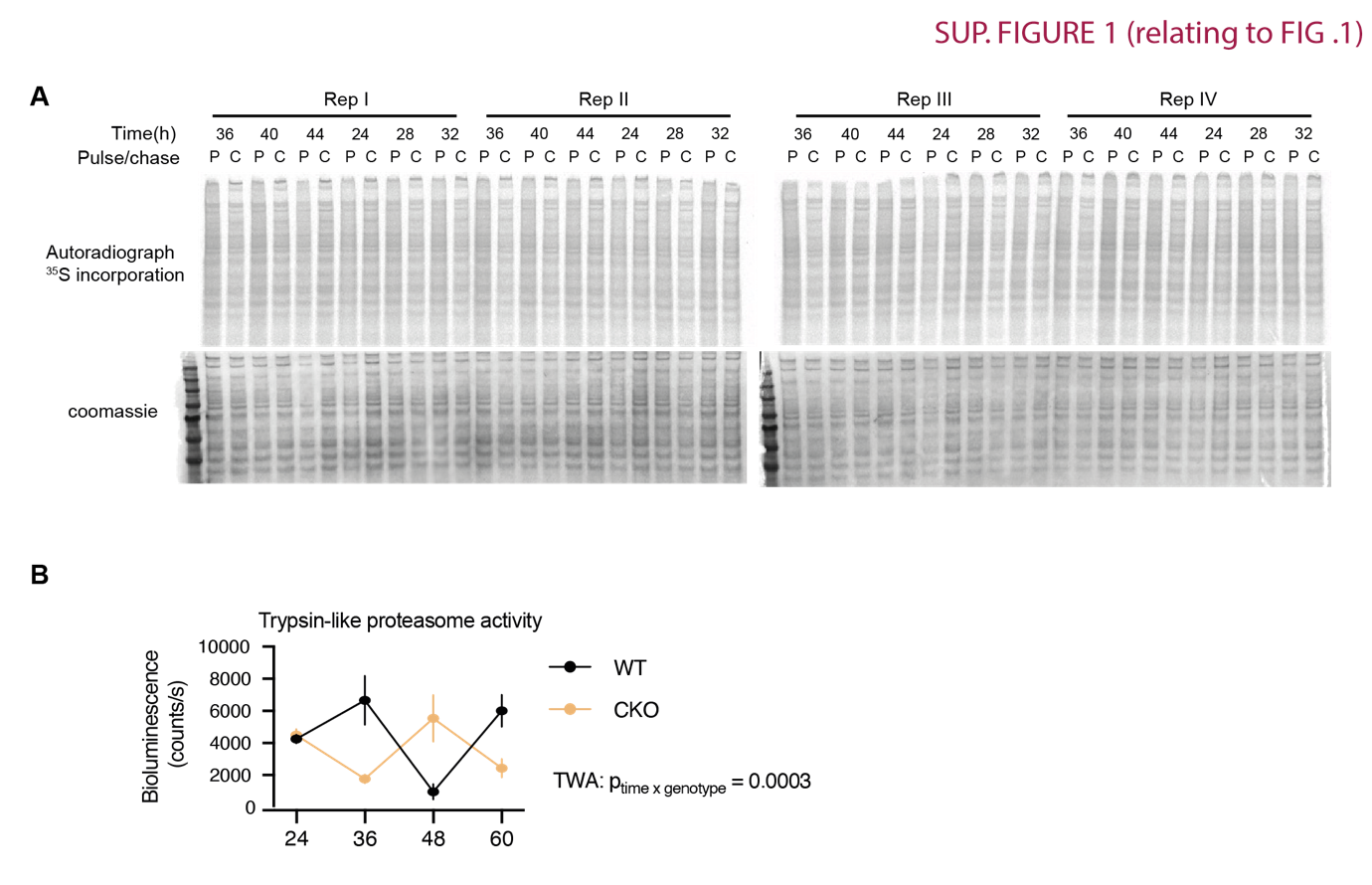

**Supplementary figure 1.**

a. Full autoradiograph and corresponding gel Coomassie stain for the ^35^S incorporation timecourse presented in Fig 1A: ^35^S-Met/Cys incorporation in 15 min pulse (P) and 1 hour chase (C) samples at different circadian times in mouse lung fibroblasts.
b. Trypsin-like proteasome activity in wild-type (WT) and *Cry1/2^-/-^* double-knockout (CKO) mouse lung fibroblasts, as measured by ProteasomeGlo cell-based assay, at different circadian times as indicated. Statistics: two-way ANOVA, p-value for interaction between the genotype and time is displayed; mean +/- SEM, n=6.

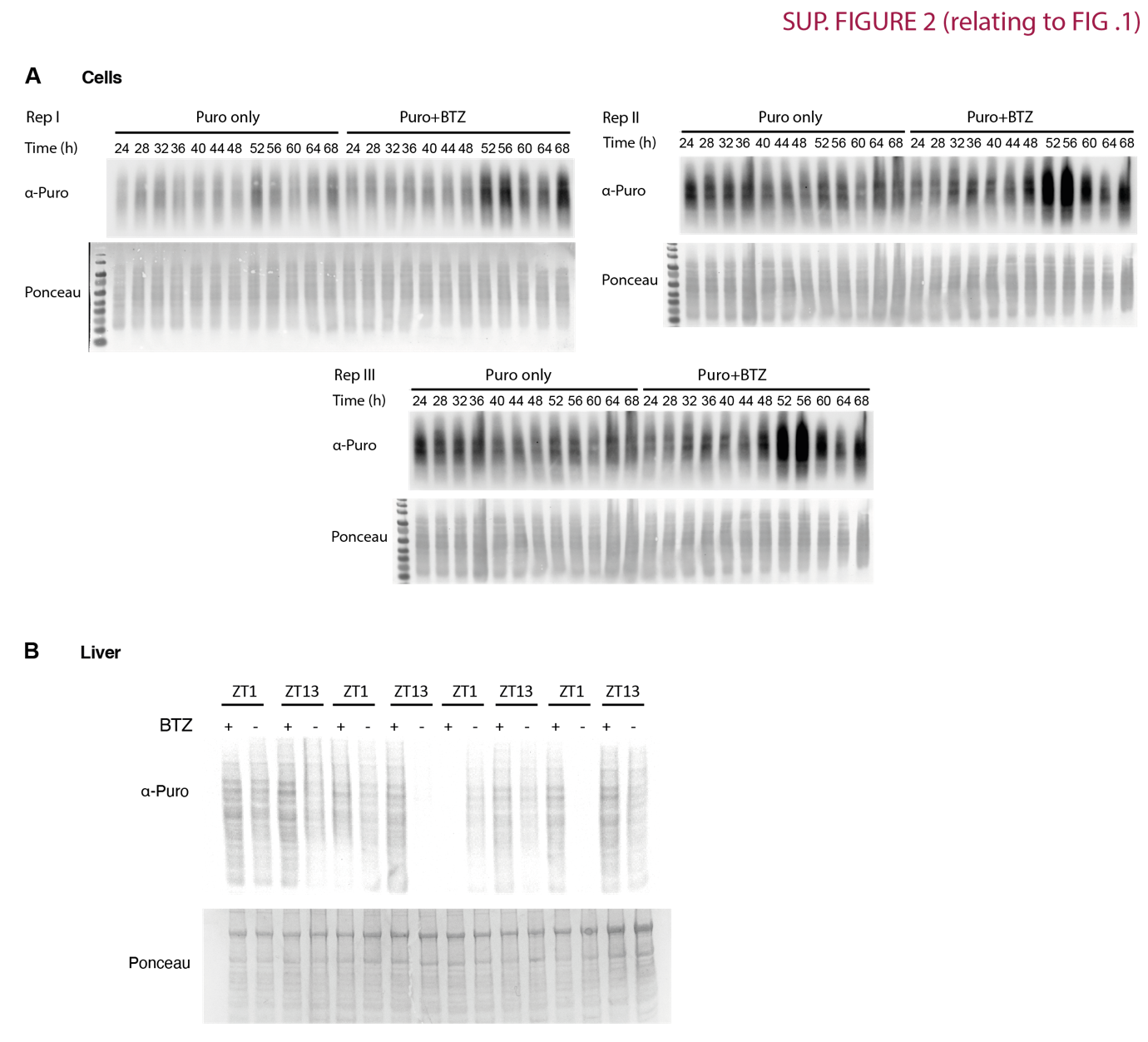

**Supplementary figure 2.**

a. Full anti-puromycin western blot and corresponding membrane Ponceau Red stain for puromycin incorporation timecourse, with 3 replicates, as also presented in Fig 1E: at each timepoint puromycin (Puro) with or without bortezomib (BTZ) was added directly to cell media, and cells lysed 30 min afterwards. Note loading order was different in replicate 3, but this does not affect quantification.
b. Full anti-puromycin western blot and corresponding membrane Ponceau Red stain for puromycin incorporation ± BTZ in mouse liver. 4 replicates, representative also presented in Fig 1F.

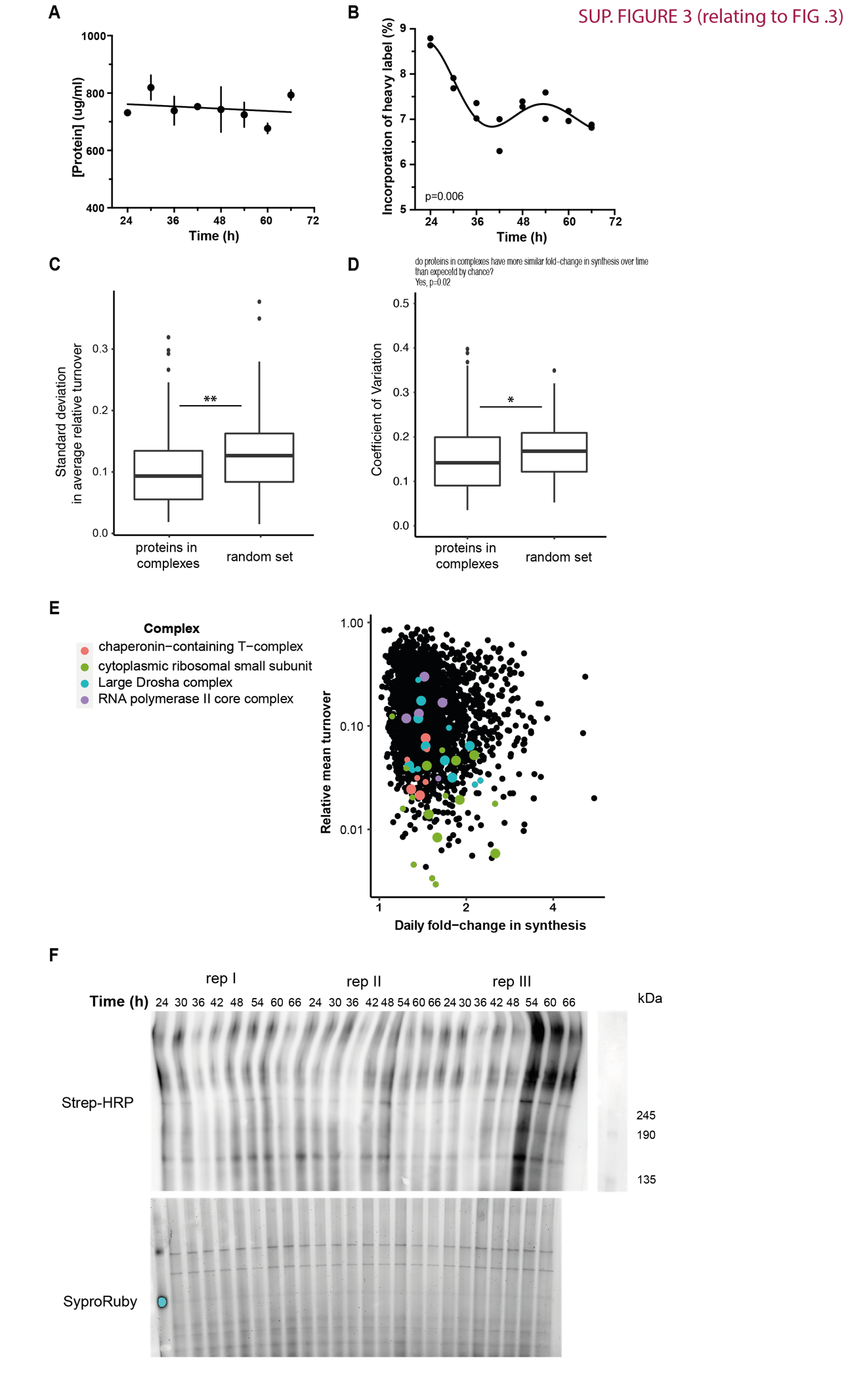

**Supplementary figure 3.**

a. Protein concentration in macromolecular complex (MMC) fraction across the pulsed SILAC timecourse, as measured by BCA assay. Statistics: straight line (null hypothesis) preferred over damped cosine wave fit, extra sum-of-squares F test p>0.05.
b. Incorporation of heavy label over the timecourse, quantified as summed intensity of heavy peptides over summed intensity of both heavy and light peptides in each sample. Only peptides detected in both heavy and light forms were considered (belonging to 2302 peptides analysed here). Statistics: damped cosine wave fit compared with straight line (null hypothesis) by extra sum-of-squares F test, the statistically preferred fit is plotted & p-value displayed.
c. Variation in relative turnover (proportion of heavy to total peptide intensity averaged across 8 timepoints, y axis), expressed as standard deviation, between proteins belonging to annotated complexes with at least 3 members, compared to a random set of proteins, randomly grouped to match the number of members in annotated complexes. Statistics: Mann-Whitney test, p<0.01.
d. Variation in protein synthesis change over circadian time (fold-change between peak and trough) expressed as coefficient of variation, between proteins belonging to annotated complexes with at least 3 members, compared to a random set of proteins, randomly grouped to match the number of members in annotated complexes. Statistics: Mann-Whitney test, p<0.05.
e. Relating to Fig 3F, coordinated turnover of proteins belonging to complexes: for four selected complexes, their annotated subunits (according to a compilation of CORUM, COMPLEAT and manual annotations) are shown individually, plotted in terms of fold-change in their synthesis over time (x-axis), and relative turnover (proportion of heavy to total peptide intensity averaged across 8 timepoints, y axis). Points that are bigger in size denote protein subunits that were significantly rhythmic (RAIN p<0.05) in their synthesis.
f. (Top) Full Native-PAGE Strep-HRP western blot of all replicates, for AHA incorporation timecourse presented in Fig 3H. (Bottom) A parallel gel was run for quantification of total protein using SyproRuby stain.

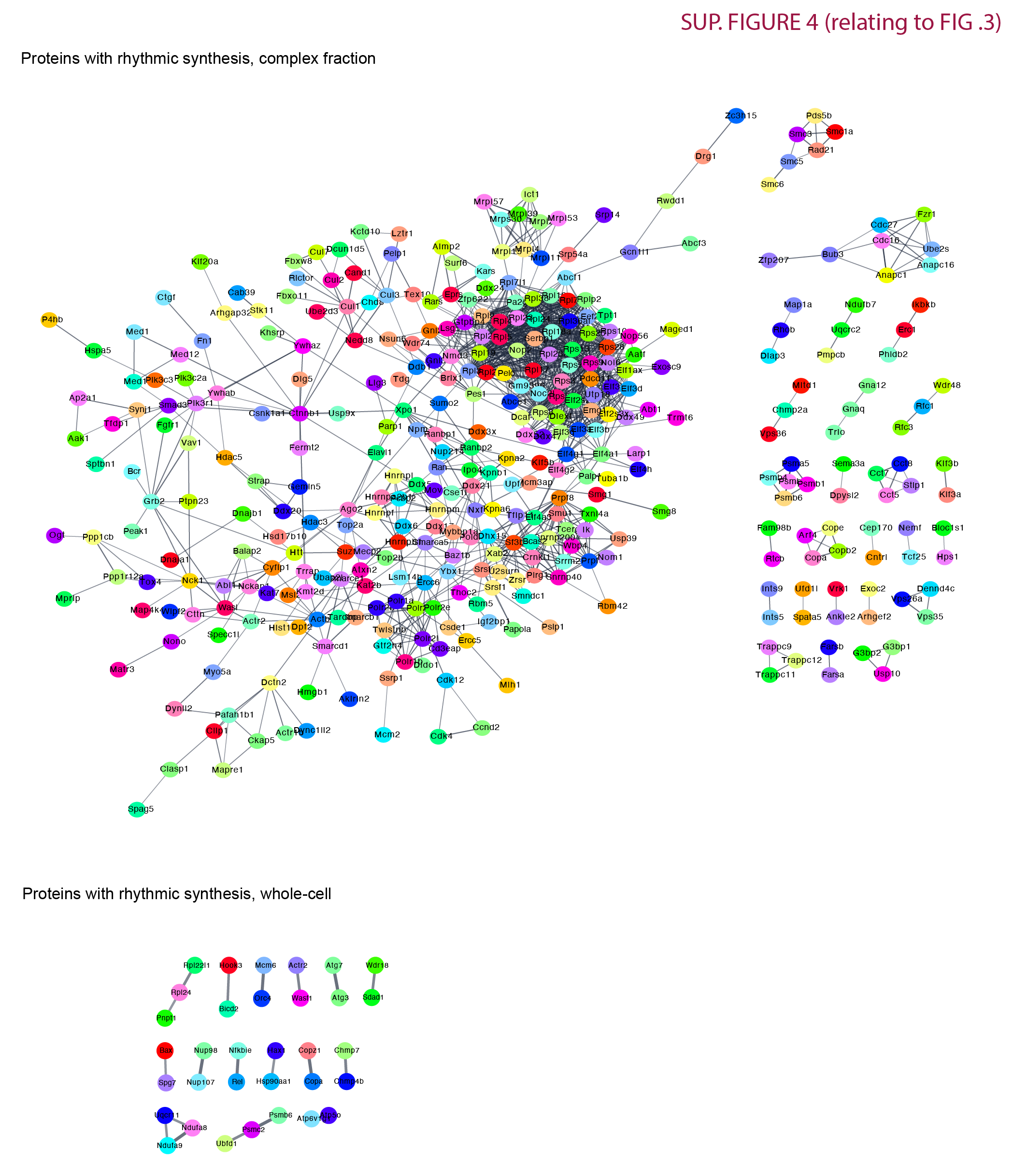

**Supplementary figure 4.**

STRING interaction network, displaying high confidence, annotated physical interactions between proteins rhythmic (RAIN p<0.05) in their synthesis in MMC fraction. Proteins with rhythmic synthesis in the complex fraction had a large, interconnected protein-protein interaction network, with an average node degree greater than 4 and a significant enrichment in interactions over all detected proteins in that experiment (q= 4.78e-5). In the bottom right, the much smaller interaction network between proteins rhythmic in their synthesis at the whole-cell level (relating to Fig. 2) is shown for comparison (on average <1 node degree, no enrichment in PPIs, q=0.3).

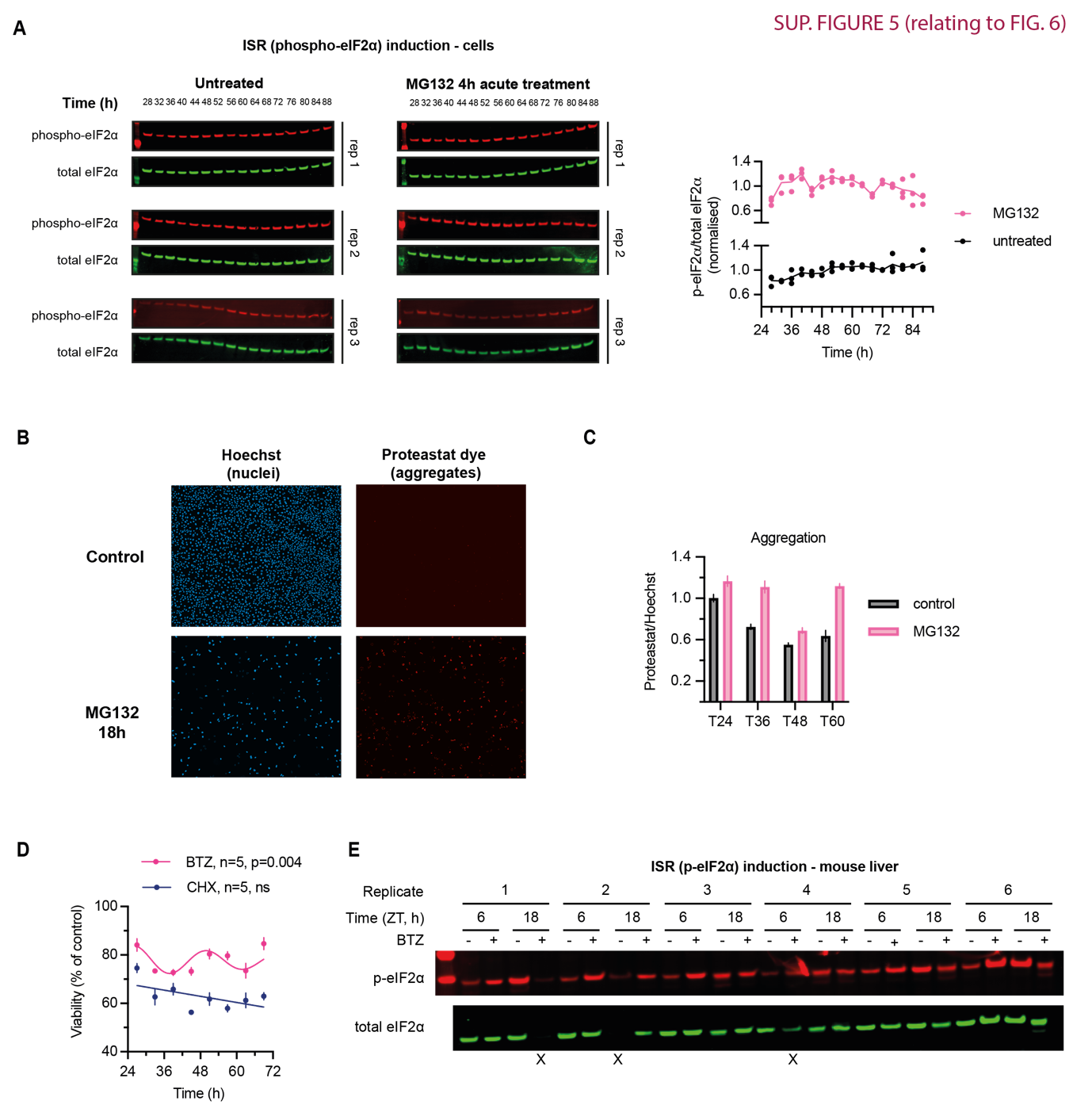

**Supplementary Figure 5.**

a. All replicate Western blots, probed for total (green) and S51-phopshorylated (red) eIF2α, of fibroblast lysates collected every 4 h for 3 days, one untreated control and one treated with 20 µM MG132 proteasomal inhibitor for 4 h before each collection.

Quantification of basal (untreated) levels of eIF2α phosphorylation (i.e. p-eIF2α/total) at each timepoint, in addition to ratio quantification in Fig 6A. Statistics: straight line fit (null hypothesis, plotted) preferred over damped cosine fit, extra sum-of-squares F test p>0.05.

b. Representative images of control fibroblasts (top row) and fibroblasts treated with MG132 overnight (bottom row), stained for nuclei with Hoechst (blue), and for protein aggregates with Proteastat molecular rotor dye (red).
c. Relative aggregation (Proteastat/Hoechst fluorescence signal) after a 24-h 20 µM MG132 or vehicle control treatment initiated at the indicated times.
d. An independent biological replicate of experiment in 5d. At 8 timepoints throughout 2 days, fibroblasts were treated with 2.5 µM proteasomal inhibitor bortezomib (BTZ), 25 µM translation inhibitor cycloheximide (CHX), or vehicle control; after 6 h, the drugs were washed out, allowing cells to recover for further 18 h. Cellular viability after the treatments, as measured by PrestoBlue High Sensitivity assay, is expressed as a proportion of control (vehicle-treated) cells at each timepoint. Statistics: damped cosine wave fit compared with straight line (null hypothesis) by extra sum-of-squares F test, the statistically preferred fit is plotted & p-value displayed.
e. Full Western blot of mouse liver lysates, probed for total (green) and S51-phopshorylated (red) eIF2α, corresponding to Fig 6E and F. Three samples were excluded, marked with X, due to undetectable or outlier levels of eIF2α.
